# Sequence-to-function embeddings across individuals capture rare and common variant heritability

**DOI:** 10.64898/2026.11.23.768687

**Authors:** Federico Billeci, Miquel Anglada-Girotto, Alexander Gusev, Jonathan Frazer, Mafalda Dias

**Author notes:** these authors contributed equally (Federico Billeci, Miquel Anglada-Girotto).

## Abstract

Statistical genetics learns how DNA shapes gene expression from variation across individuals, whereas sequence-to-function models learn from variation along the genome. Because sequence-to-function models map sequence to molecular readouts through deep architectures, their intermediate representations could offer function-informed descriptions of genetic variation. Whether these representations capture differences between individuals, however, is unknown. Here we introduce Embedding-based Heritability (EMBHER), a variance-component framework that estimates the expression heritability captured by these representations. Applied to Borzoi, embeddings recover much of the whole-blood heritability explained by genotypes and far exceed the model’s final predictions. Deeper layers increasingly prioritize functional, evolutionarily constrained sequence while encoding less ancestry information than genotypes. Embeddings also explain more rare-variant heritability than genotypes, consistent with sharing information across variants too rare to estimate individually. EMBHER integrates sequence-to-function models with statistical genetics, a first step towards population-to-function models.

---

Two largely separate approaches seek to explain how DNA sequence shapes gene expression. Statistical genetics learns from variation across individuals: expression quantitative trait locus (eQTL) mapping, transcriptome-wide association studies (TWAS) and heritability analysis relate genotypes to expression differences in population cohorts^1–3^. Current sequence-to-function (S2F) models, on the other hand, learn from variation along the genome: deep neural networks trained on the reference genome and functional genomics data predict gene expression, chromatin accessibility, and transcription factor binding from sequence windows of up to a megabase^4–6^.

Learning from different axes of variation, the two approaches are likely to capture complementary aspects of how genotype shapes expression. Conventional statistical genetics approaches represent genetic variation through genotype dosage matrices, which do not explicitly encode variant function; incorporating functional annotations can enable information sharing between functionally similar variants and improve power for rare-variant association testing^7^. Such information sharing is particularly valuable for rare and ultra-rare variants, which are sparsely represented across individuals in population cohorts despite collectively accounting for a substantial fraction of the cis-heritability of gene expression^8^. Moreover, these approaches typically assume additive variant effects, limiting their ability to capture how the effect of a variant depends on its haplotype context or how multiple variants interact non-additively.

S2F models are widely used to predict variant effects by comparing predictions for reference and alternative sequences. They have shown promising performance in predicting the effects of individual common and rare variants across diverse molecular readouts, including gene expression, splicing, and chromatin accessibility, as evaluated using molecular QTL data and high-throughput functional assays, including MPRAs and saturation mutagenesis^4,5,6,9–13^. They could therefore, in principle, capture the effects of rare variants and singletons and their combinations within individual haplotypes. However, when applied to personal genomes from population cohorts, S2F models perform substantially worse at predicting inter-individual differences in gene expression than statistical genetics approaches trained on genotype dosages^14–17^. Fine-tuning S2F models on paired genomic and transcriptomic data improves predictions for genes seen during training, but generalization to unseen genes remains limited^16^.

As deep neural networks, S2F models progressively transform input DNA sequences into intermediate representations, or embeddings, before generating their final predictions. These embeddings should provide a function-informed representation of genetic variation, and may retain information about inter-individual expression variation that the final predictions do not reflect. In this letter, we explore this idea and its implications for the scope of S2F models. Quantifying the inter-individual expression information calls for a measure that does not depend on training a new predictor on limited data. Heritability provides such a measure, as it upper-bounds the held-out variance explained by any linear predictor built on the same representation^1,2^. We introduce Embedding-based Heritability (EMBHER), a variance-component framework^18–20^ that estimates the cis-expression heritability attributable to S2F embeddings by replacing the genetic relatedness matrix with an embedding-based relatedness matrix (Fig. 1).

**Figure 1.**
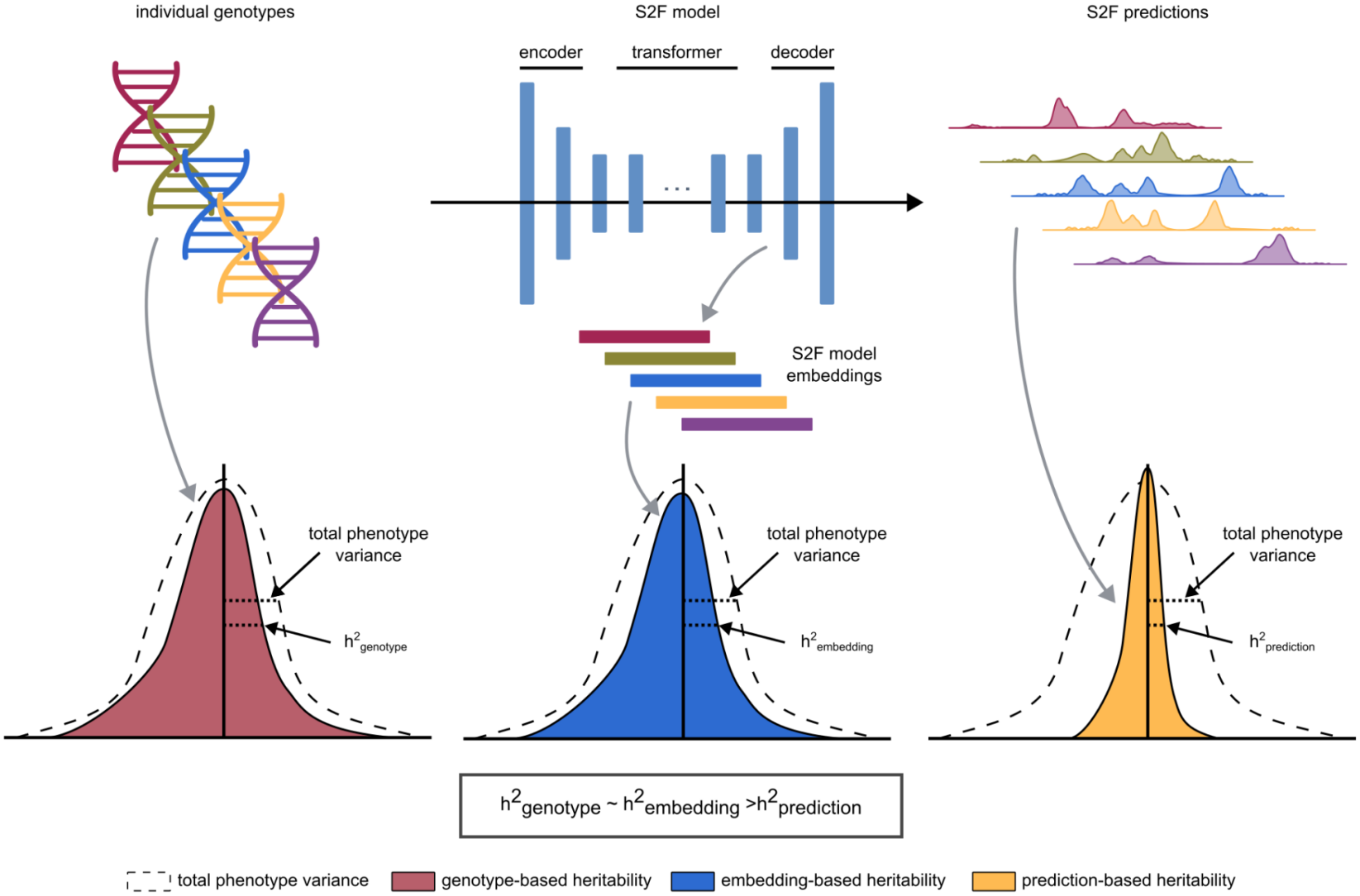
EMBHER estimates the expression heritability captured by sequence-to-function (S2F) model embeddings. Schematic of Embedding-based Heritability (EMBHER). Individual genotypes are passed through a S2F model to obtain intermediate embeddings and final molecular predictions. For each representation, the proportion of phenotypic variance explained (h^2^, coloured areas) is compared with the total phenotypic variance (dashed curve): genotype-based (h^2^_G_, blue), embedding-based (h^2^_E_, red) and prediction-based (h^2^_P_, green) heritability.

EMBHER extends conventional genotype-based heritability estimation by replacing the genetic relatedness matrix with an embedding-based relatedness matrix, allowing the estimation of the proportion of expression variance explained by the embeddings: the *embedding-based heritability* (h^2^_E_). To account for the high dimensionality and correlation structure of neural network embeddings, EMBHER incorporates dimensionality reduction and whitening (Supplementary Note 1). We validated the approach through extensive simulations across different genetic architectures and embedding properties (Supplementary Note 2; Supplementary Fig. 1-4).

## S2F embeddings capture cis-expression heritability

We applied EMBHER to Borzoi, a 190-million-parameter S2F model, using whole-blood RNA-seq and genomic data from 800 GTEx donors. For each gene and donor, we obtained embeddings for 24 intermediate layers as well as whole-blood expression predictions by passing the personalized sequence window centred on the transcription start site (TSS) through Borzoi (Methods; Supplementary Table 1). We computed genotype-based heritability (h^2^_G_) on all the genes with TPM>0.1 in at least 50% of whole-blood GTEx donors (Methods; Supplementary Table 2). We analysed 500 randomly selected genes with h^2^_G_>0.01 in whole blood. For each gene, we compared the proportion of expression variance captured by embeddings (h^2^_E_) with conventional genotype-based heritability, as a baseline (Supplementary Table 3). Consistent with previous benchmarks^15–17,21–23^, Borzoi’s whole-blood predictions recover substantially less expression heritability (h^2^_P_) than conventional genotype-based estimates (Fig. 2a, left). However, in sharp contrast, embeddings from the penultimate layer of the same model recover much of this gap, capturing substantially more expression variance than the final model predictions (Fig. 2a, right). These findings are consistent with those obtained in the independent GEUVADIS cohort, comprising 449 donors with whole-genome sequencing and lymphoblastoid cell-line RNA-seq (Supplementary Fig. 5). Together, these findings suggest that S2F models learn representations that capture inter-individual expression variation without requiring fine-tuning, even when their predictions fail to reflect this information.

**Figure 2.**
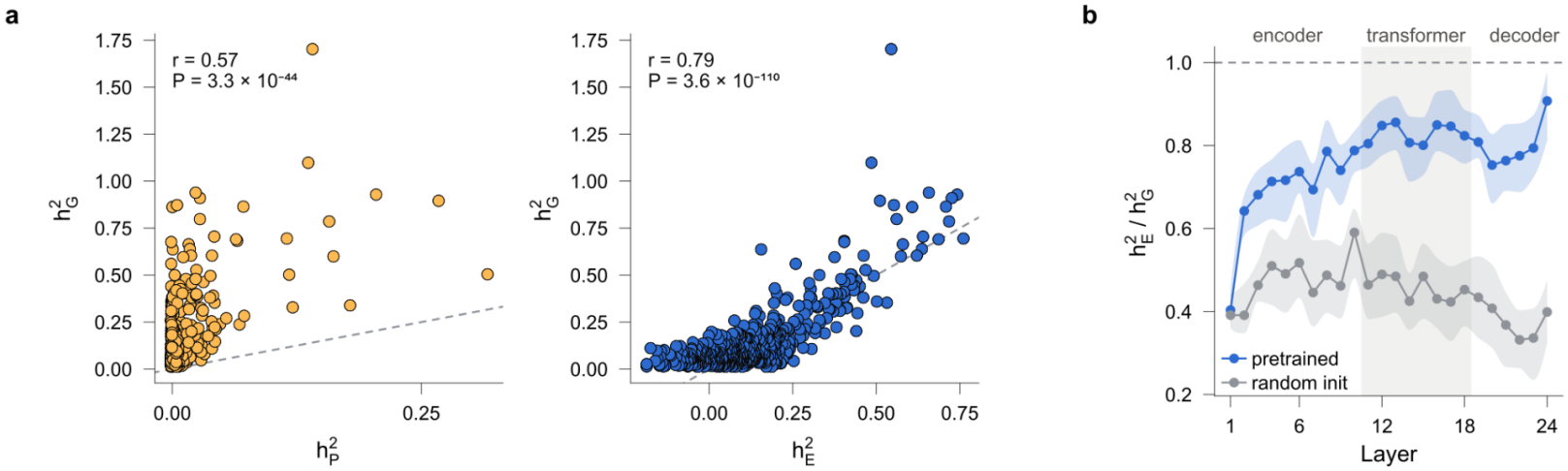
Sequence-to-function (S2F) model embeddings capture inter-individual expression variation that final predictions do not reflect. Analyses use GTEx whole blood (n = 800 donors) and 500 randomly selected genes with h^2^_G_ > 0.01. (a) Genotype-based expression heritability (h^2^_G_) plotted against the heritability explained by Borzoi’s whole-blood predictions (h^2^_P_, left) and by embeddings from the penultimate layer (h^2^_E_, right). Each point represents a gene; dashed lines indicate equality (y = x); r, Pearson correlation. (b) Median ratio of embedding-based to genotype-based heritability (h^2^_E_/h^2^_G_) across successive Borzoi layers for the pretrained (red) and randomly initialized (grey) model. Shaded regions indicate 95% bootstrap confidence intervals across genes. The dashed line at a ratio of 1 indicates heritability equal to the genotype-based estimate.

Embedding-based heritability increases across successive Borzoi layers, approaching genotype-based heritability in deeper layers (Fig. 2b). This increase is not observed in a randomly initialised Borzoi model, suggesting that it arises from training rather than from the model architecture alone. Furthermore, while we find information on inter-individual variation largely absent from single tissue-specific predictions, it is recovered when RNA-seq predictions across all GTEx tissues are concatenated (Supplementary Fig. 6). This suggests that information relevant to inter-individual expression variation is distributed across tissue-specific outputs, rather than fully captured by individual tracks, which may not fully reflect the information learned by the model.

## S2F embeddings capture functional signals beyond population structure

The above observations raise the question of whether embeddings merely recapitulate genotype-based similarity or capture additional, functionally meaningful relationships between sequences that conventional genotype dosage matrices overlook. We assessed the extent to which each representation captures local ancestry. We trained linear models to predict European versus African ancestry using the first ten principal components of each Borzoi embedding layer, and compared their performance with analogous models trained on genotype principal components from the same genomic window (Methods; Supplementary Table 4). Across all layers, embeddings explained substantially less ancestry-associated variation than genotypes (Fig. 3a, left; Supplementary Fig. 7). This pattern is illustrated for a randomly chosen gene in Fig. 3a (right): the first two principal components of the gene-locus genotype matrix show substantially greater separation between ancestry groups than those of the embedding matrix. These results imply that embeddings are less influenced by population structure than genotype dosages.

**Figure 3.**
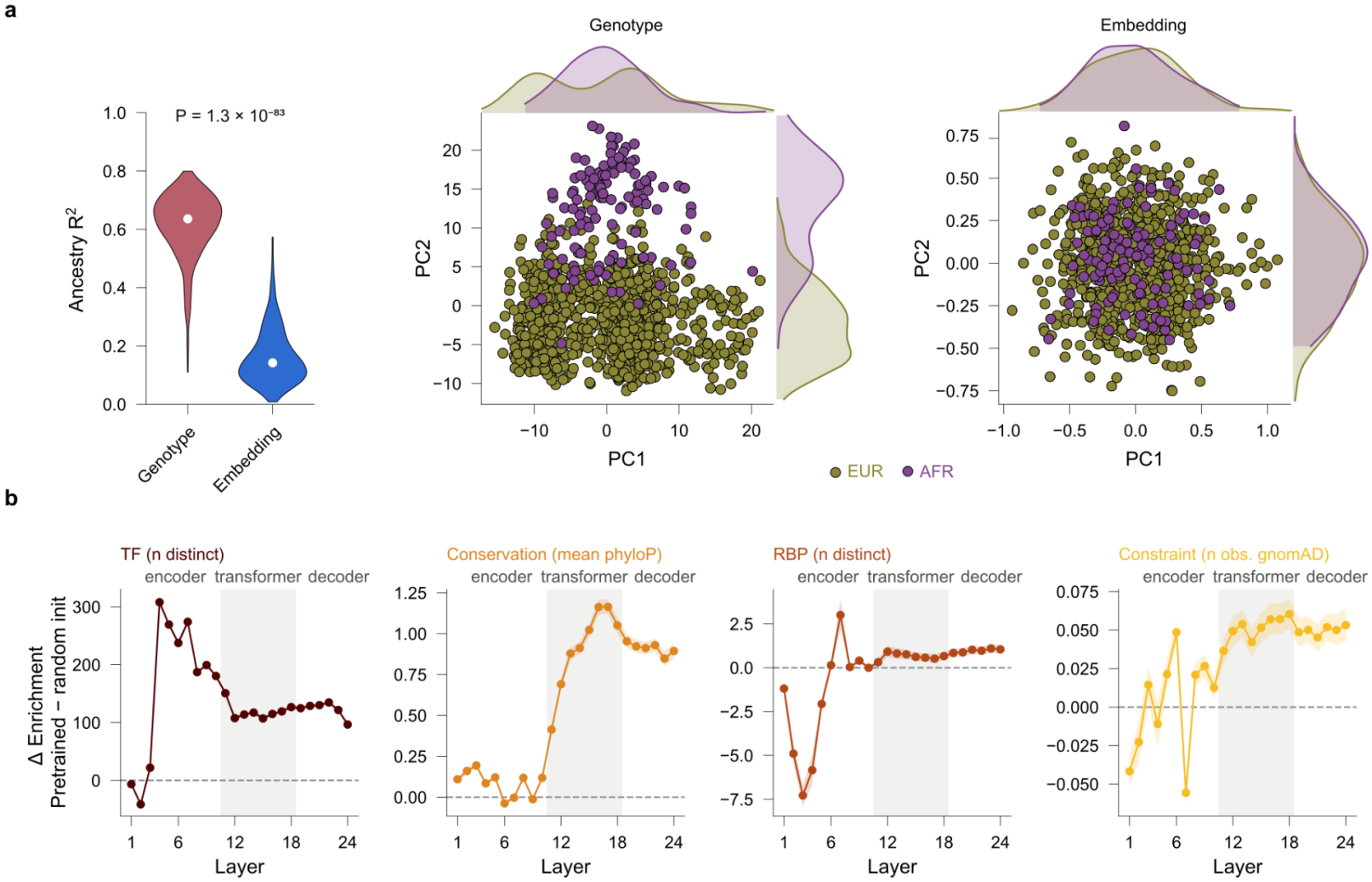
Embeddings are less influenced by population structure and progressively prioritize functionally relevant sequence features. (a) Ancestry separability of genotypes and Borzoi embeddings. For each of 500 genes, R^2^ of a linear model predicting ancestry (EUR, European, n = 800; AFR, African, n = 122) from the top 10 principal components (PCs) of the genotype matrix or of the final Borzoi embedding of the gene’s 524-kb window. R^2^ is computed in-sample and measures separability, not prediction accuracy. Violins show the distribution across genes (white dot, median; black bar, interquartile range); P, two-sided paired Wilcoxon signed-rank test across genes. Scatter plots show PC1 and PC2 of the genotype matrix (left) and embedding matrix (right) for an example gene (FAAP20), with donors coloured by ancestry and marginal densities per group. The layer-wise analysis is shown in Supplementary Fig. 7. (b) Difference in sequence-attribution metrics between the pretrained and randomly initialized Borzoi models across layers (positive values indicate greater enrichment in the pretrained model). Metrics, from top left: enrichment of unique RNA-binding protein motifs (CisBP-RNA), enrichment of unique transcription factor motifs (HOCOMOCO), evolutionary conservation (phyloP), and enrichment of variants unobserved versus observed in gnomAD.

## S2F embeddings progressively prioritize functionally relevant sequence features

To investigate which sequence features contribute to these representations, we applied input x gradient attribution^24^ to identify highly contributing sequence regions (seqlets) for each gene and Borzoi layer (Fig 3b; Supplementary Table 5). We computed enrichments of functional annotations within and outside seqlets. To distinguish learned functional features from effects arising solely from nonlinear sequence transformations, we compare these enrichments with a randomly initialised Borzoi model. Constraint and conservation per gene (as measured by mean phyloP and observed in gnomAD) are increasingly enriched in later layers, whereas molecular interaction annotations such as the number of unique transcription factors (TF) are especially recognised in early layers, stabilising in later layers. A similar, while noisier, picture arises for RNA-binding protein (RBP) motifs. These results suggest that S2F models progressively develop representations that prioritize diverse biologically relevant sequence features, rather than merely transforming genotype information through nonlinear operations. Hence, by organizing genetic variation according to learned functional properties, embeddings may reveal similarities between individuals that are overlooked by conventional genotype-based representations.

## S2F embeddings expose rare-variant heritability

Embeddings not only contain different information but also reveal more about the effect of genetic variation than could be captured by genetic association alone. In particular, in the case of rare variants where there is limited statistical power, we find that the functionally-informed representation improves genotype-based estimates of interindividual variation (Fig. 4). We extended our analysis to all high-quality genes, partitioned genetic variation by allele frequency and generated embeddings exclusively from rare variants (MAF ≤ 0.01), comparing their heritability estimates (h^2^_E,rare_) with those obtained from the corresponding rare-variant genotypes (h^2^_G,rare_; Methods; Supplementary Fig. 8). Embedding-based heritability estimates significantly exceed genotype-based estimates (bootstrap p < 0.002; Fig. 4a; Supplementary Fig. 8). This suggests that S2F embeddings capture functionally meaningful inter-individual variation attributable to rare variants by better reflecting phenotypic distance between individuals (Fig. 4b).

**Figure 4.**
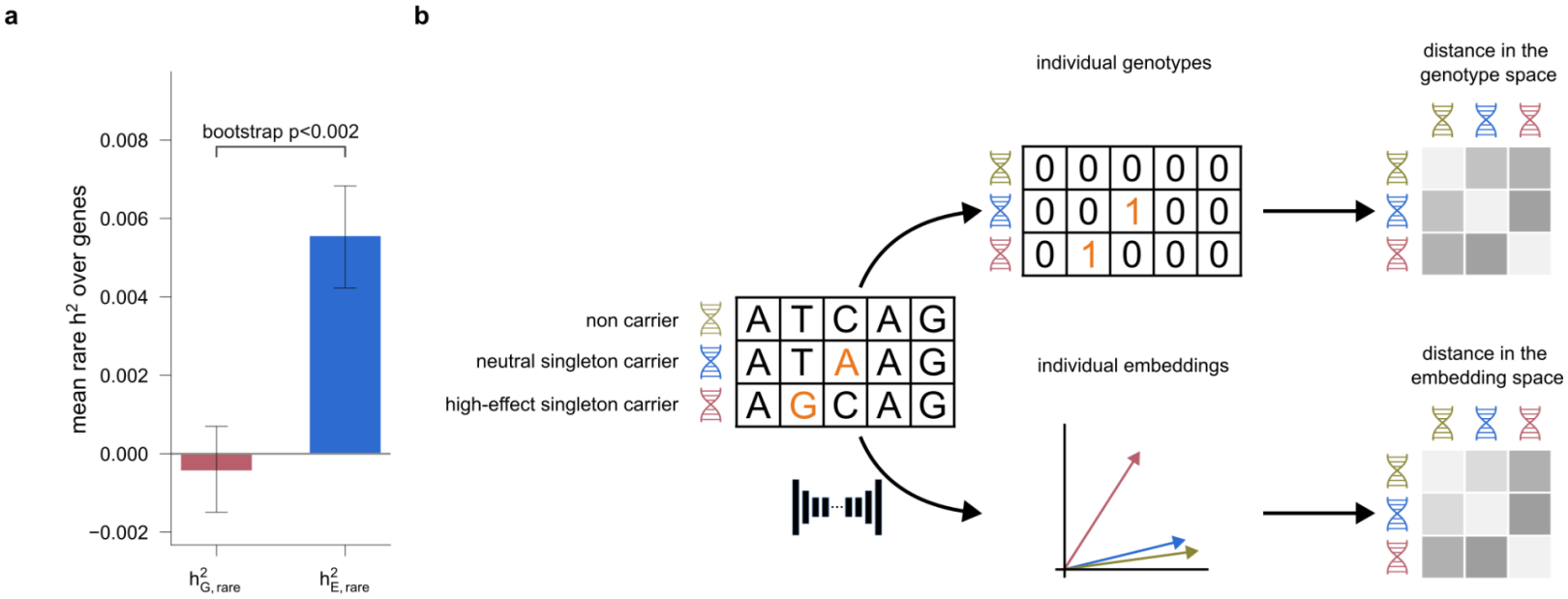
Function-informed embeddings capture rare-variant-associated expression heritability. (a) Transcriptome-wide mean heritability of rare variants (minor allele frequency, MAF ≤ 0.01) estimated from genotypes (h^2^_G, rare_) and from embeddings (h^2^_E, rare_) across all high-quality genes in GTEx whole blood. Error bars indicate 95% confidence intervals from bootstrapping genes; bootstrap P < 0.002. (b) Schematic illustrating how a neutral and a high-effect singleton variant can yield similar genotype-based relatedness but distinct embedding-based relatedness between individuals.

Our results demonstrate that S2F models trained on the reference genome learn intermediate representations that capture a substantial fraction of inter-individual expression variation, even when this information is not reflected in their final predictions. The poor performance of S2F models on personal genomes therefore appears to reflect, at least in part, how this information is extracted by the output head rather than what the models have learned. This distinction has implications both for how S2F models are evaluated and improved, and for how statistical genetics can make use of them.

For S2F models, EMBHER enables population-scale benchmarking of learned representations by estimating the expression heritability they capture, independently of any particular output track. As the field moves from single-variant to population-scale evaluation, embedding-based heritability provides a measure of how much inter-individual variation a model’s representations contain, independently of how well its output head extracts it. Tracking this quantity across layers also reveals where information about inter-individual variation is gained or lost within the network, potentially informing the design of model architectures, training strategies and loss functions. Finally, because heritability upper-bounds the held-out variance explained by linear predictors, embedding-based heritability indicates the maximum performance attainable by a linear model trained on frozen embeddings. Such models offer a lightweight alternative to fine-tuning, which is computationally costly and so far generalizes poorly to genes not seen during training^16^. Personalized predictions would then require only forward passes through the frozen S2F model and a set of linear weights for each gene.

For statistical genetics, S2F embeddings provide information complementary to the genotype dosages used in most methods, and their value is likely to be greatest for rare variants. Frequency-based approaches to estimating variant effects are fundamentally limited by the vast number of possible variants in the genome relative to the size of any realistic cohort. Recent population growth has produced a vast excess of rare variants, many of them private to a single family or individual^25,26^, so even sequencing the entire human population would leave most variants with too few carriers for their effects to be estimated. In cohorts with matched whole-genome sequencing and molecular phenotypes, which rarely exceed a few thousand donors, the limit applies already at minor allele frequencies around 1%. Function-informed representations can, in principle, sidestep this limit by sharing information between variants with similar functional effects, however rarely each is observed. Consistent with this, embeddings captured more rare-variant heritability than genotypes. Incorporating embedding-based relatedness into heritability partitioning, transcriptome-wide association studies and rare-variant association tests would test this directly.

Although this study focused on bulk RNA-seq, EMBHER can be applied to any phenotype measured in a cohort, not only to the molecular readouts on which the S2F model was trained. Because S2F embeddings integrate regulatory features learned across many molecular readouts, embedding-based relatedness could also be applied to complex traits and diseases. Comparing the heritability captured by different layers, or by predictions for different tissues and assays, could then indicate which regulatory features and cell types are most relevant to each trait.

S2F embeddings are nonlinear functions of each haplotype’s sequence and can therefore, in principle, capture interactions between variants and the dependence of a variant’s effect on its haplotype context. Quantifying the expression variance attributable to such effects is a natural next step, with implications for rare pathogenic variants. Many of these variants show reduced penetrance in population cohorts compared with family-based studies^27,28^, and regulatory variation on the same haplotype is one plausible contributor^29^. Moving from the effects of isolated variants to the effects of whole haplotypes could therefore help explain why carriers of the same pathogenic variant differ in disease risk.

These findings come with limitations. The analyses used a single S2F model, Borzoi, and two cell types, whole blood and lymphoblastoid cell lines. Embeddings were averaged across sequence positions and summed across both haplotypes, which may discard information, particularly about rare variants carried on a single haplotype. Applying EMBHER to other S2F models, such as AlphaGenome, and to other tissues will show how general these findings are.

Our results provide a proof of principle that deep learning models of sequence function can be integrated into the variance-component framework of statistical genetics. Combining what S2F models learn along the genome with what statistical genetics learns across individuals is a first step towards population-to-function models.

## METHODS

Detailed descriptions of the simulation experiments and assessments of robustness to model assumptions are provided in the Supplementary Notes accompanying this manuscript. Here, we describe the datasets, computational procedures, and statistical methods used in the main analyses.

### GTEx data

We analyzed whole-blood gene expression and phased whole-genome sequencing (WGS) data from 800 donors in the GTEx project. WGS genotypes were aligned to GRCh38 and phased using SHAPEIT2. Gene expression data were obtained from the GTEx v10 gene-level transcripts per million (TPM) matrix. We selected the 803 whole-blood RNA-seq samples included in the analysis freeze, retained the first sample for donors with multiple samples, and matched these donors to those with available phased WGS data, yielding 800 individuals.

Gene expression values were normalized in two steps. First, expression values were quantile-normalized across genes within each donor to the mean distribution across donors, averaging ranks in the case of ties. Second, a rank-based inverse-normal transformation was applied to each gene across donors.

Covariate adjustment was performed within the heritability models rather than directly on the normalized expression values. We used the covariates provided for GTEx v10 whole-blood eQTL mapping (https://gtexportal.org/home/methods), including 60 PEER factors, the first five genotype principal components, sequencing platform, PCR protocol, and sex.

### GEUVADIS data

For independent validation, we analyzed gene expression and phased whole-genome sequencing (WGS) data from 449 donors in the GEUVADIS lymphoblastoid cell-line (LCL) cohort. WGS genotypes were obtained from the 1000 Genomes Project 30× high-coverage dataset, comprising 3,202 individuals, with variants aligned to GRCh38.

Gene expression data were obtained from the GEUVADIS GD660 gene-level RPKM matrix, containing 660 sequencing runs. RPKM values were converted to transcripts per million (TPM) for each run, and technical replicates were averaged to obtain one expression profile per donor. We retained donors with available phased WGS data, yielding 449 individuals. Gene expression values were normalized using the same two-step procedure applied to GTEx.

As for GTEx, covariate adjustment was performed within the heritability models. We used a comparable set of covariates, including 60 PEER factors, the first five genotype principal components, sex, and sequencing centre. PEER factors were calculated from log_2_(TPM + 1) values using genes with TPM > 0 in at least 50% of donors. Genotype principal components were obtained from genomi-ancestry-panel (github.com/exon-research/genomi-ancestry-panel), which provides precomputed principal components for the 1000 Genomes 30× dataset.

### High-quality gene selection

High-quality genes were required to have TPM >0.1 in at least 50% of donors, yielding 19,845 and 20,320 genes for GTEx whole blood and GEUVADIS lymphoblastoid cell lines, respectively. We retained only autosomal genes (chromosomes 1–22) annotated in GENCODE v46. To ensure compatibility with the Borzoi input window, we selected only genes whose entire gene body lies within ±97,304 bp of the transcription start site, as Borzoi predicts only the central 196,608 bp of its full 524,288-bp input window. The final gene sets contained 16,671 GTEx genes and 15,861 GEUVADIS genes. The analyses in Figs. 2–3 and Supplementary Fig. 5 were performed on a subset of 500 genes with h^2^_G_ > 0.01 to compare methods in the presence of a stronger genetic signal.

### Sequence-to-function modeling with Borzoi

We used the replicate 0 of the PyTorch version of Borzoi^5^ with its published pretrained weights, implemented using the borzoi-pytorch port. Model parameters were kept frozen throughout the analyses. All forward passes were performed on GPUs using bfloat16 autocasting.

#### Personalized sequences

We adapted the personal genome evaluation framework from Tu et al.^15^. For each gene, we extracted a 524,288-bp genomic window centred on its TSS using the GRCh38 primary assembly and GENCODE v46 annotations. We reconstructed the two haplotype-specific sequences for each donor by applying all phased variants within the corresponding window to the reference genome. Because insertions and deletions alter sequence length, reconstructed haplotypes were centre-trimmed to the required 524,288-bp input length. Ambiguous nucleotides (N) were encoded as all-zero vectors. Each haplotype was processed independently, requiring two forward passes per donor and gene.

#### Embeddings

We extracted intermediate activations from 24 layers spanning the Borzoi architecture: the initial convolution (*encoder_conv_dna*), five residual convolutional blocks (*encoder_res_tower_0-4*), U-net skip convolutions (*skip_horizontal_conv0-1*) and bottleneck (*encoder_unet1, encoder_trunk*), eight transformer blocks (*transformer_block_0-7*), decoder upsampling and separable-convolution stages (*decoder_upsample1, decoder_separable1, decoder_upsample0, decoder_separable0*), the cropped trunk output (*embeddings_cropped*), and the final convolution preceding the output head (*embeddings_final*).

Each layer produced an activation tensor with dimension channels by positions. We averaged activations across sequence positions to obtain one vector per haplotype, then summed the two haplotype vectors to obtain a single embedding per donor, genomic window and layer. Unless otherwise specified, analyses used the final-layer embedding (*embeddings_final*), consisting of 1,920 features obtained after a 1×1 convolution and GELU activation. Embeddings represent the entire genomic window centered on the gene TSS rather than a gene-specific prediction.

#### Predicted expression

We extracted predicted RNA-seq coverage from the three GTEx whole-blood output tracks (indices 7531, 7532 and 7533). Borzoi’s output squashing transformation was inverted using its original parameters (soft-clipping threshold 384, scale 0.01 and exponent 3/4). For each gene and haplotype, we summed the resulting coverage across prediction bins overlapping the union of exons annotated in GENCODE v46. Predictions were then summed across the two haplotypes and averaged across the three whole-blood tracks on the linear scale, yielding one predicted expression value per donor and gene.

#### Predicted expression along other tissues

To assess how much inter-individual variation in measured whole-blood expression is captured by predictions across tissues, we used all 1,543 Borzoi RNA-seq output tracks, including both strands of stranded assays. For each gene, haplotype, and track, we inverted the output squashing transformation using the published track-specific scale, soft-clipping threshold (384) and exponent (3/4), and summed the resulting coverage across prediction bins overlapping the union of exons annotated in GENCODE v46. Predictions were then summed across haplotypes on the linear scale, yielding a 1,543-dimensional vector per donor and gene. These vectors were treated as embeddings, and heritability was estimated using the same embedding-based relatedness-matrix construction and Haseman-Elston regression framework applied to intermediate Borzoi embeddings, as described below. Because this representation was obtained from the prediction head rather than an intermediate layer, it was evaluated separately from the layer-wise analysis (Supplementary Fig. 6).

#### Randomly initialized control

To distinguish the contribution of pretraining from effects arising from the architecture alone, we generated a randomly initialized Borzoi model with the same architecture, configuration, and output head as the pretrained model. Parameters were initialized using PyTorch defaults with a fixed random seed (0). A single random initialization was used, and embedding extraction, heritability estimation, and attribution analyses were repeated on the same genes as for the pretrained model.

### Heritability estimation

We estimated the proportion of gene expression variance explained by genotype dosages, Borzoi embeddings, and Borzoi expression predictions using a linear mixed model variance-component framework. For each gene, we fitted the model

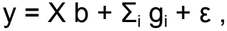

where g_i_ ~ N(0, σ^2^_i_K_i_), ε ~ N(0, σ^2^_e_I), y is the preprocessed gene expression phenotype, X contains an intercept and covariates as fixed effects, and K_i_ is the relatedness matrix for component i. For GTEx, fixed effects included 60 PEER factors, the first five genotype principal components, sequencing platform, PCR protocol, and sex. For GEUVADIS, fixed effects included 60 PEER factors, the first five genotype principal components, sex, and sequencing centre.

Variance components were estimated using multi-component Haseman-Elston (HE) regression^30,31^. The estimator is unconstrained, and negative estimates were retained. The heritability of each component was calculated as its contribution to the total estimated variance. Analyses were performed using EMBHER, with further methodological details and validation provided in Supplementary Notes 1-2.

#### Genotype-based heritability (h^2^ _G_)

For each gene, we extracted all SNVs and indels within the 524,288-bp genomic window. Genotypes were standardized and partitioned into six bins based on minor allele count (MAC), minor allele frequency (MAF), and linkage disequilibrium (LD): MAC ≤ 5; MAC > 5 and MAF ≤ 0.01; MAF in (0.01, 0.1], split into low- and high-LD bins; and MAF in (0.1, 0.5], similarly split by LD. High or low LD was defined as the bottom or top 50% of LD scores in the gene. For each bin, we constructed a genetic relatedness matrix K = ZZ^T^/M, where Z is the standardized genotype matrix, and M is the number of variants. All six components were fitted jointly, and h^2^_G_ was defined as the sum of their estimated heritabilities.

#### Embedding-based heritability (h^2^_E_)

For each gene and Borzoi layer, we constructed a relatedness matrix from the donor × feature embedding matrix. To account for correlations between embedding features, we standardized the features and applied singular value decomposition. We retained the first 70% of the numerical rank and whitened the resulting components before constructing the relatedness matrix K = WW^T^/k, where W contains the whitened components and k is their number. This procedure was applied consistently across all layers. Details of the whitening procedure and its statistical properties are provided in Supplementary Note 1.

#### Rare-variant heritability

We compared genotype- and embedding-based heritability attributable to rare variants (MAF ≤ 0.01) across all 16,671 high-quality GTEx genes. Genotype-based rare-variant heritability (h^2^_G,rare_) was defined as the sum of the first two components of the six-bin genotype model. For embedding-based rare-variant heritability (h^2^_E,rare_), the first two components were replaced by a single rare-variant embedding kernel, while the four common-variant components were retained.

Rare-variant embeddings were generated by applying only variants with MAF ≤ 0.01 to each donor’s haplotype sequences, keeping all other positions at the reference allele. Embeddings were prepared and whitened as described above. Both models used identical phenotypes, donors, and covariates.

#### Prediction-based heritability (h^2^_P_)

For each gene, Borzoi-predicted expression was standardized across donors and used to construct a rank-one relatedness matrix. Heritability was estimated using the same HE estimator and covariates as in the genotype- and embedding-based analyses. Because the predictions form a one-dimensional vector, with one value per donor, the variance explained could alternatively be quantified by the squared Pearson correlation between covariate-adjusted predictions and observed expression (partial R^2^), yielding practically identical estimates. We retained the HE approach for consistency across analyses.

#### Layer-wise analysis

We estimated h^2^_E_ for each of the 24 Borzoi layers and calculated the ratio h^2^_E_ /h^2^_G_ for each gene. Results were summarized by the median ratio across genes, with 95% confidence intervals obtained from 2,000 bootstrap resamples. Pretrained and randomly initialized models were evaluated on the same genes.

### Ancestry prediction

We assessed how strongly genotypes and Borzoi embeddings distinguish population groups using 922 GTEx donors self-reporting as White (EUR, n = 800) or Black or African American (AFR, n = 122). Embeddings were computed from the full 944-donor WGS cohort. For each of the 500 selected genes, we extracted genotype dosages within the 524,288-bp window and embeddings from each Borzoi layer.

For each representation, we performed principal component analysis (PCA) and fitted a linear regression of the binary population group label (EUR = 0, AFR = 1) on the first 10 principal components. Group separability was measured using the coefficient of determination (R^2^). For the layer-wise analysis, results were summarized by the median ratio of embedding-based to genotype-based R^2^, with 95% bootstrap confidence intervals.

### Attribution-based enrichment analysis

#### Genes, sequences and models

We analysed the 500 randomly selected genes. For each gene, the input was the reference sequence (GRCh38) of the 524,288-bp window centred on the TSS, without individual genetic variation. Every analysis was run on pretrained Borzoi and on a randomly initialized control with the same architecture and output head, so that the effect of training is read as the difference between the two.

#### Attribution and seqlets

For each of the 24 layers used for the embeddings, the attribution target was the L2 norm of the position-averaged activation vector. We computed input×gradient^24^ of each target with respect to the one-hot input and summed across the four nucleotide channels to obtain a per-position score. We called seqlets (4-25 bp) on each score leveraging the Tangermeme framework’s recursive seqlet caller with *p* < 5·10^-4^, once per gene, model and layer. All annotation analyses below use these seqlets.

#### Transcription factor (TF) and RNA-binding protein (RBP) motifs

We scanned a 36-bp window centred on each seqlet with the FIMO algorithm^32^ as implemented in the gReLU package at *p* < 1·10^-4^ against HOCOMOCO v13 CORE^33^ (TFs, both strands) or CisBP-RNA^34^ (RBPs, sense strand only, restricted to seqlets within the gene span, with minus-strand genes reverse-complemented). Motifs were collapsed to TFs or RBPs, counting each once per window. Because FIMO finds chance hits at this threshold, we drew 20,000 random 36-bp windows per gene from the same sequence (or gene span for RBPs), scanned them identically, and used the fraction carrying each TF or RBP as its null rate. For each motif carried by at least two seqlet windows, a one-sided binomial test compared the number of seqlet windows carrying it with the null expectation; p-values were adjusted per gene and layer with the Benjamini-Hochberg procedure, and motifs with q < 0.05 were called enriched. The readout is the number of enriched TFs or RBPs per gene and layer. Because RBP motif analysis was restricted to seqlets within gene coordinates, the number of seqlets tested varied substantially with gene length. We therefore adjusted RBP counts, but not TF counts (which were obtained from the full sequence window), to the mean log number of seqlets using a per-layer regression slope estimated across genes after centring within each model and shared between pretrained and randomly initialized models.

#### Evolutionary conservation and population constraint

Each position in the window was assigned to one of 32 trinucleotide contexts (reverse-complement collapsed) and classified as seqlet or background (all other positions, with nothing excluded). For conservation, we scored positions with the 241-way mammalian phyloP track (Zoonomia) and computed ΔphyloP as the mean difference between observed phyloP at seqlet positions and the expectation given their context, defined as the mean phyloP of background positions of the same context, pooled over genes. For population constraint, a position was labelled seen if gnomAD v4.1.1 genomes contain a PASS SNV with allele count > 0 there, and unseen otherwise; positions with low coverage were excluded. We computed the log odds ratio of unseen versus seen positions in seqlets relative to the context-matched expectation, where positive values indicate depletion of variation. Both statistics were pooled over genes.

### Permutation test and bootstrap p-values for rare variant heritability analysis

We performed two complementary analyses of rare-variant heritability across genes. First, to quantify uncertainty in the across-gene estimates, we generated 1,000 paired bootstrap resamples of genes, sampling with replacement and retaining both model estimates for each sampled gene. Within each resample, we calculated the mean heritability and paired difference between models (Δh^2^), and obtained 95% percentile confidence intervals (Fig. 4b).

To assess whether the across-gene mean and median Δh^2^ = h^2^_E,rare_ − h^2^_G,rare_ exceeded expectations under a no-association null, we generated 1,000 independent donor-label permutations of each gene’s expression phenotype, holding kernels and covariates fixed. Both models used identical permutations within each gene, and each permutation replicate yielded an across-gene mean and median Δh^2^. Two-sided permutation P-values were calculated as (1 + number of null statistics with absolute magnitude at least as large as observed)/1,001. Results of the permutation tests are illustrated in Supplementary Fig. 8.

### Software and computational implementation

Heritability estimation was performed using our implementation in EMBHER. Simulation experiments were adapted from the publicly available Simulator repository (https://github.com/sriramlab/Simulator)^35^. Genotype processing and LD-score calculations used PLINK 2^36^, and genomic windows were extracted using bcftools^37^. Borzoi inference and embedding extraction were implemented in PyTorch using borzoi-pytorch^38^. Attribution analyses used Captum^39^, Tangermeme^40^, Decima^41^, and gReLU^42^.

Computational pipelines were executed using Nextflow 24.10^43^ and Snakemake^44^ on a SLURM-managed computing cluster.

## DATA AVAILABILITY

### Intermediate files generated from analyzing data analyses

Intermediate files generated in this study, including the genomic coordinates (GRCh38/hg38) of Borzoi attribution seqlets identified across 500 genes, are publicly available on Zenodo (https://doi.org/10.5281/zenodo.24246557).

Individual-level GTEx controlled-access data and restricted derivatives are not redistributed with this manuscript.

### Previously published datasets used

#### GTEx

Data from the Genotype-Tissue Expression (GTEx) Project were obtained from the NIH database of Genotypes and Phenotypes (dbGaP; study accession phs000424) and the GTEx Portal (https://gtexportal.org/home/downloads/adult-gtex).

Controlled-access data included phased whole-genome sequencing (WGS) genotypes aligned to GRCh38 (GTEx_Analysis_2021-02-11_v9_WholeGenomeSeq_944Indiv_Analysis_Freeze.SHAP EIT2_phased.vcf.gz; dbGaP accession phg001796.v1) and individual-level subject phenotypes (phs000424.v11.pht002742.v9.p2.c1.GTEx_Subject_Phenotypes.GRU.txt.gz), used to define population groups in the ancestry analyses. Access to these data requires authorization through dbGaP.

Publicly available GTEx Analysis v10 data were downloaded from the GTEx Portal, including gene-level expression estimates (GTEx_Analysis_v10_RNASeQCv2.4.2_gene_tpm.gct.gz), sample and subject annotations (GTEx_Analysis_v10_Annotations_SampleAttributesDS.txt and GTEx_Analysis_v10_Annotations_SubjectPhenotypesDS.txt), and tissue-specific eQTL covariates (GTEx_Analysis_v10_eQTL_covariates.tar).

### GEUVADIS and 1000 Genomes Project

GEUVADIS gene expression data were obtained from ArrayExpress/BioStudies (accession E-GEUV-1; https://www.ebi.ac.uk/biostudies/arrayexpress/studies/E-GEUV-1). Gene-level expression estimates were obtained from GD660.GeneQuantRPKM.txt.gz, and sample metadata, including sex and sequencing centre, from E-GEUV-1.sdrf.txt.

Phased genotypes were obtained from the 1000 Genomes Project 30× high-coverage resequencing panel (3,202 individuals), using the GRCh38 phased SNV/indel/structural-variant VCFs (1kGP_high_coverage_Illumina.chr{1-22}.filtered.SNV_INDEL_SV_phased_panel.vcf.g z), available at https://ftp.1000genomes.ebi.ac.uk/vol1/ftp/data_collections/1000G_2504_high_coverage/working/20220422_3202_phased_SNV_INDEL_SV/. The first five genotype principal components were obtained from genomi-ancestry-panel v1.0.0 (https://github.com/exon-research/genomi-ancestry-panel).

### Reference genome and annotations

The GRCh38 reference genome and GENCODE v46 gene annotations were obtained from https://ftp.ebi.ac.uk/pub/databases/gencode/Gencode_human/release_46/.

Transcription factor binding motifs were obtained from HOCOMOCO v13 CORE (https://hocomoco13.autosome.org/downloads_v13), and human RNA-binding protein motifs from CisBP-RNA (http://cisbp-rna.ccbr.utoronto.ca; accessed 14 October 2025). Evolutionary conservation scores were obtained from the Zoonomia 241-way mammalian phyloP track (cactus241way.phyloP.bw), available through the UCSC Genome Browser (https://hgdownload.soe.ucsc.edu/goldenPath/hg38/cactus241way/). Population variation annotations were obtained from gnomAD v4.1.1 genome sites VCFs (https://storage.googleapis.com/gcp-public-data--gnomad/release/4.1.1/vcf/genomes/).

### Model weights

Pretrained Borzoi replicate 0 model weights were obtained from Hugging Face (https://huggingface.co/johahi/borzoi-replicate-0).

## CODE AVAILABILITY

The full data analysis pipeline of this manuscript is publicly available at https://github.com/TheBil99/EMBHER_publication and the corresponding standalone package to run EMBHER at https://github.com/TheBil99/EMBHER.

## ACKNOWLEDGEMENTS

We thank Sílvia Bonàs-Guarch, Donate Weghorn and Jonathan Pritchard for their valuable feedback and insightful discussions during the development of this work; Andrew Marderstein and Joshua Popp for the helpful discussion on the interpretation of Borzoi embeddings; Elena Galioto for the help with plots. We thank the participants and donors of the Genotype-Tissue Expression (GTEx) project, the GEUVADIS study, and the 1000 Genomes Project for their invaluable contributions, and the respective consortia for generating and making these data available to the scientific community. We also acknowledge the support of the Spanish Ministry of Science and Innovation through the Centro de Excelencia Severo Ochoa (CEX2020-001049-S, MCIN/AEI/10.13039/501100011033), the Generalitat de Catalunya through the CERCA programme and the EMBL partnership, andAnthropic, through the AI for Science program. Research for this publication has been partially carried out in the Barcelona Collaboratorium for Modelling and Predictive Biology.

## AUTHOR CONTRIBUTIONS

FB and MA-G conceived and designed the study; FB developed the statistical methodology, the theoretical framework and the simulations; MA-G adapted the deep-learning methods and developed the attribution analysis; FB and MA-G cleaned and analyzed the data; AG contributed to the statistical methods development and simulation studies; FB, MA-G, AG, JF and MD interpreted the results; FB, MA-G, JF and MD wrote the manuscript; JF and MD supervised the project and acquired the funding. All authors read and approved the final manuscript.

## Supplementary Materials

### 1 Supplementary Note 1

#### 1.1 The EMBHER model

EMBHER is a linear mixed-model framework to estimate the proportion of inter-individual phenotypic variance captured by representations learned by sequence-to-function (S2F) models. EMBHER extends genotype-based heritability estimation by replacing the genetic relatedness matrix with a relatedness matrix constructed from individuals’ S2F embeddings.

In a genotype-based linear mixed model, expression levels for a given gene across *n* individuals are modeled as:

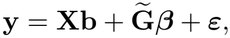

where:

- **y** *∈* ℝ^*n*^ is the vector of normalized gene expression levels across individuals.
- **X** *∈* ℝ^*n×p*^ is the design matrix of *p* fixed-effect covariates, with coefficients **b** *∈* ℝ^*p*^.
- 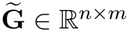 is the genotype dosage matrix of *m* variants, centered and standardized across individuals.
- ***β*** *∈* ℝ^*m*^ contains the effect sizes associated with each variant.
- ***ε*** *∈* ℝ^*n*^ represents residual variation.

We assume that the genetic effect sizes are independent and identically distributed Gaussian random variables:

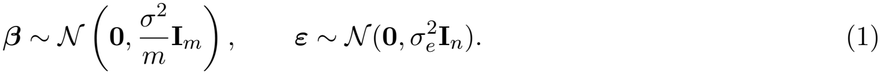

Assuming independence between genetic effects and residual variation, the resulting phenotype covariance matrix is:

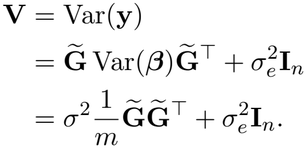

Defining the genetic relatedness matrix (GRM) as:

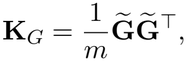

the covariance matrix can be written as:

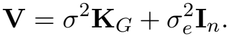

EMBHER generalizes this framework by replacing the standardized genotype matrix with a matrix of S2F embeddings. After dimensionality reduction and whitening, the resulting embedding matrix 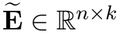 is used in the analogous linear mixed model. The construction of 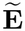 from the original S2F embeddings is described in the following section.

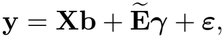

where ***γ*** *∈* ℝ^*k*^ contains the effects associated with the retained embedding components. Analogously to Eq. (1), we assume that these effects are independent and identically distributed Gaussian random variables:

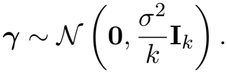

The corresponding phenotype covariance matrix is:

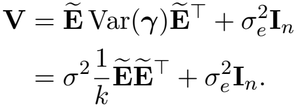

Defining the embedding-based relatedness matrix (ERM) as:

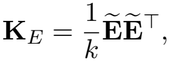

we obtain:

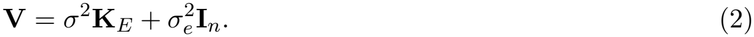

Thus, EMBHER uses the standard linear mixed-model variance-component formulation, with a relatedness matrix constructed from S2F embeddings.

For either representation, the proportion of phenotypic variance attributable to the relatedness matrix is defined as:

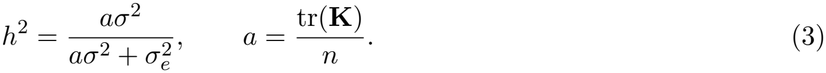

Here, **K** denotes either the genetic or embedding-based relatedness matrix, and *a* is its mean diagonal, accounting for the scale of the relatedness matrix. This definition applies independently of the method used to estimate the variance components.

##### 1.1.1 Dimensionality reduction and whitening

For each gene, we start from an embedding matrix **E** *∈* ℝ^*n×d*^, where *n* is the number of individuals and *d* is the number of embedding features. We first standardize each feature across individuals to have zero mean and unit variance, obtaining the matrix **E**_*s*_.

We then perform a compact singular value decomposition (SVD) of the standardized embedding matrix:

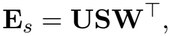

where **U** *∈* ℝ ^*n×r*^ contains the left singular vectors, **S** *∈* ℝ ^*r×r*^ is a diagonal matrix of nonzero singular values ordered from largest to smallest, and **W** *∈* ℝ^*d×r*^ contains the right singular vectors. Here, *r ≤* min(*n−*1, *d*) is the numerical rank of **E**_*s*_, namely the number of principal components retained after excluding singular values with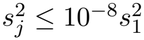, where *s*_1_ is the largest singular value. Zero-variance features are removed before the SVD; feature standardization uses population standard deviations.

We retain the first *k ≤ r* principal components using a truncation matrix **T**_*k*_ *∈* ℝ^*r×k*^, defined as:

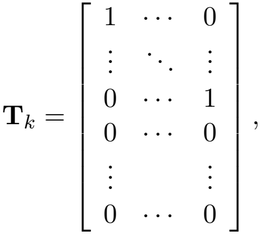

where the first *k* rows form an identity matrix and the remaining *r − k* rows contain zeros. Multiplication by **T**_*k*_ selects the first *k* columns of **U**, corresponding to the principal components with the largest singular values.

We choose *k* = round(0.7*r*), retaining 70% of the available principal components (rounded to the nearest integer).

We construct the truncated and whitened embedding matrix as:

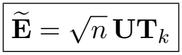

where 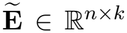. This transformation is equivalent to projecting the standardized embeddings onto their first *k* principal components and dividing each component by its root-mean-square magnitude across individuals.

Dimensionality reduction is important for variance-component estimation because retaining all *n −* 1 nonzero principal components, when the standardized embedding matrix has maximum rank *r* = *n −* 1, yields:

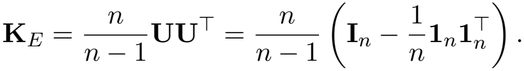

This matrix has unit diagonal and constant off-diagonal entries equal to *−*1*/*(*n −* 1), independently of the original embedding representation. After accounting for the intercept, the resulting covariance structure is proportional to the identity matrix, making the embedding-associated and residual variance components non-identifiable. We therefore retain only a subset of principal components to preserve informative differences in the relatedness structure between individuals.

##### 1.1.2 Embedding-based relatedness matrix

The ERM is calculated from the truncated and whitened embeddings as:

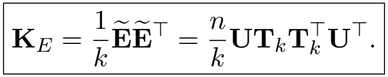

Each element (**K**_*E*_)_*ij*_ represents the similarity between individuals *i* and *j* across the retained embedding components. The resulting ERM can therefore replace the conventional GRM in the variance-component model defined in Eq. (2).

This construction is related to Mahalanobis relationship matrices previously proposed to account for linkage disequilibrium in genotype-based heritability estimation ^1,2^. These approaches incorporate inverse genotype covariance to account for correlations among SNPs. Here, we apply an analogous principle to embedding features through empirical whitening restricted to the retained principal components.

#### 1.2 Fitting the EMBHER model

EMBHER has two estimator branches: EMBHER-HE (the default), which uses Haseman–Elston moment regression, and EMBHER-REML, which uses restricted maximum likelihood. Both branches fit the variance-component model defined in (2). Both estimators internally use the standardized phenotype 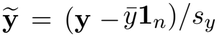, where *s*_*y*_ is the phenotype standard deviation

##### 1.2.1 EMBHER-HE

In EMBHER-HE, we estimate the variance components by Haseman–Elston (HE) regression using an all-entry moment-regression formulation.

We first fit the fixed effects by ordinary least squares. For a full-column-rank design matrix **X**, define:

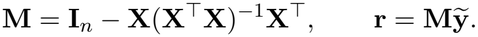

Writing **A**_1_ = **K** and **A**_2_ = **I**_*n*_, the residual covariance is:

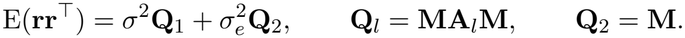

EMBHER-HE fits these moments by least squares over all *n × n* matrix entries, including the diagonal and both symmetric off-diagonal entries:

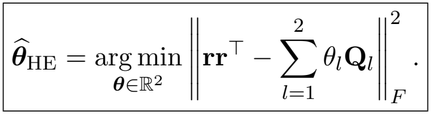

Here ∥ · ∥_*F*_ denotes the Frobenius norm and 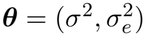. Equivalently, let **D** have columns vec(**Q**_1_) and vec(**Q**_2_), where vec stacks all matrix entries, and set **t** = vec(**rr**^⊤^). The normal equations are:

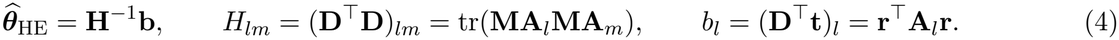

###### Standard errors

Uncertainty in EMBHER-HE heritability is estimated by a delete-one individual jack-knife. For each individual *i*, we remove the corresponding entries of the phenotype and covariate matrix and the corresponding row and column of each relatedness matrix. We then rebuild the covariate residual projection for the remaining *n −* 1 individuals and apply the same EMBHER-HE estimator and heritability definition, obtaining 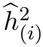. Each fold uses the mean diagonal of its retained kernel:

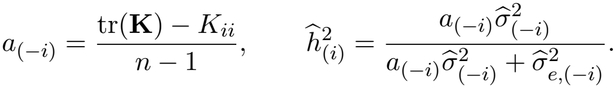

With 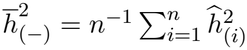, the reported standard error is:

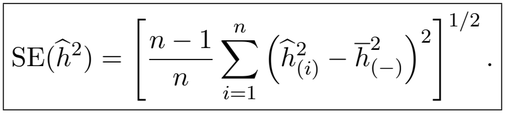

This corresponds to (*n −* 1) times the population variance of the delete-one estimates before taking the square root. The implementation obtains the fold-specific normal equations from full-sample quantities using rank-one updates to the covariate cross-products and row/column deletions of the kernels. This is algebraically equivalent to explicit refitting with covariate residualization in each fold.

##### 1.2.2 EMBHER-REML

In EMBHER-REML, we estimate variance components by restricted maximum likelihood (REML), using the same objective for genotype-based and embedding-based relatedness matrices.

Given a relatedness matrix **K**, we define:

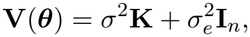

where 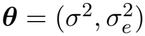 denotes the variance components to be estimated. The restricted log-likelihood is:

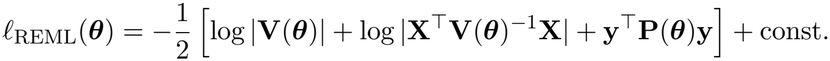

where:

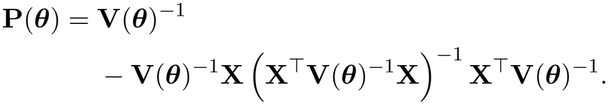

We estimate the variance components by maximizing the restricted log-likelihood over the feasible parameter space:

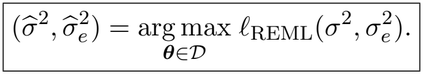

Here the feasible parameter space is:

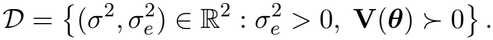

#### 1.3 Extension to multiple variance components

The framework can be straightforwardly extended to jointly estimate the variance explained by multiple relatedness matrices, constructed from either genotypes or S2F embeddings. Given *q* relatedness matrices **K**_1_, …, **K**_*q*_, the phenotype covariance matrix becomes:

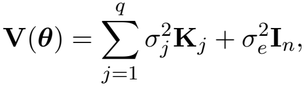

where 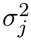 represents the variance component associated with the *j*-th relatedness matrix. The variance components are jointly estimated using the same EMBHER-REML objective defined above. Each kernel coefficient 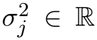 may be negative, while 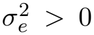 and **V**(***θ***) *≻* 0 are required for a feasible likelihood evaluation. Component heritabilities and their sum are reported without truncation.

The heritability definition in (3) extends to multiple variance components, with the contribution of each component estimated as:

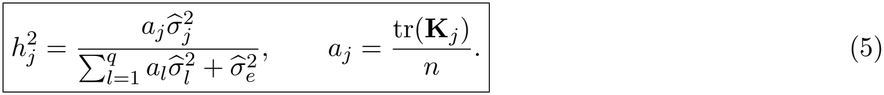

EMBHER-HE extends to the same joint model by taking **A**_*j*_ = **K**_*j*_ for *j* = 1, …, *q* and **A**_*q*+1_ = **I**_*n*_ in (4). All *q* + 1 components are estimated simultaneously by moment regression, using the same residual projection **M**. The projected component matrices must be linearly independent for the normal-equation matrix to be invertible. The same heritability definitions apply, with total heritability given by the sum of the component shares:

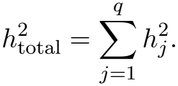

For HE jackknife standard errors, each delete-one fit uses the fold-specific mean diagonal of every kernel.

#### 1.4 Genotype-based heritability estimation

For each gene, we estimated genotype-based cis-expression heritability using all genetic variants within a 524,288-bp genomic window centered on the transcription start site (TSS). This corresponds to the same sequence window used as input to Borzoi, ensuring that genotype-based and embedding-based heritability estimates capture variation attributable to the same genomic region.

We partitioned variants into six mutually exclusive components based on minor allele count (MAC), minor allele frequency (MAF), and LD score: ultra-rare (MAC *≤* 5), rare (MAC *>* 5 and MAF *≤* 0.01), low-frequency (0.01 *<* MAF *≤* 0.1), and common (MAF *>* 0.1). Low-frequency and common variant components were each further divided into low-LD and high-LD components using the 50th percentile of the LD score distribution within each gene and frequency class.

We estimated the contributions of these components jointly using a multiple-component linear mixed model:

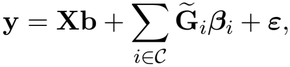

where *C* = *{*ur, r, lf_*L*_, lf_*H*_, c_*L*_, c_*H*_*}* denotes the six variant components, with subscripts *L* and *H* indicating low- and high-LD variants, respectively. Here, 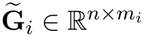 is the centered and standardized genotype matrix containing *m*_*i*_ variants from component *i*, and 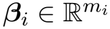 contains their effect sizes.

Under the variance-component model described above, we assume independent Gaussian-distributed genetic effects:

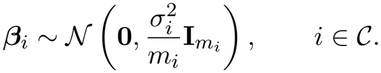

Assuming independence between components, the resulting phenotype covariance matrix is:

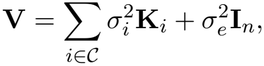

where the genetic relatedness matrix for component *i* is defined as:

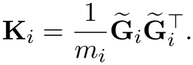

The variance components were estimated jointly using the multiple-component framework described above. The heritability associated with each component was defined as in (5). Genotype dosages are centered using the estimated allele frequency and scaled by 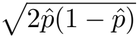.

Total genotype-based heritability was calculated as the sum of the contributions of the six components:

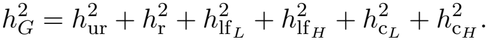

For analyses focusing on rare genetic variation, we defined rare-variant genotype-based heritability as the sum of the ultra-rare and rare components:

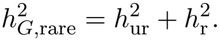

#### 1.5 Estimation of rare-variant heritability from embeddings

To estimate the contribution of rare genetic variants to inter-individual expression variation captured by S2F models, we generated embeddings from personal haplotype sequences containing only ultra-rare and rare variants, as defined above. For each individual, variants with MAF *>* 0.01 were replaced by the reference allele, while ultra-rare and rare variants were retained on their respective haplotypes.

We processed these sequences through the S2F model and constructed an embedding-based relatedness matrix **K**_*E*,rare_ using the dimensionality reduction and whitening procedure described above.

To estimate the variance associated with these embeddings while accounting for higher-frequency genetic variation, we jointly fitted **K**_*E*,rare_ with the four genotype-based relatedness matrices corresponding to the low-frequency and common variants partitioned by LD score, as defined in the previous section. The phenotype covariance matrix was:

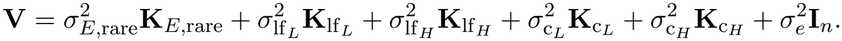

The variance components were estimated jointly using the multiple-component framework described above. We defined rare-variant embedding heritability as:

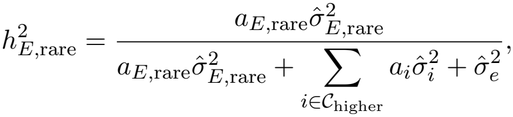

where *C*_higher_ = *{*lf_*L*_, lf_*H*_, c_*L*_, c_*H*_*}* denotes the four higher-frequency genotype components.

#### 1.6 Prediction-based heritability estimation

For each gene, we estimated prediction-based heritability using the individual-level whole-blood expression predictions obtained from Borzoi, as described in the Predicted expression section. Each individual was represented by a single predicted expression value, which was used to construct a prediction-based relatedness matrix **K**_pred_.

We then estimated the proportion of observed inter-individual expression variance explained by these predictions using a single-component variance-component model with **K**_pred_ as the relatedness matrix, following the framework described above. We denote the resulting estimate as 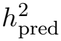, calculated as:

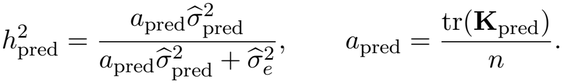

## 2 Supplementary Note 2

### 2.1 Simulation framework for genotype-based heritability estimation

Simulations were performed using real genotypes from 944 GTEx individuals, for 10 randomly selected genes. For each gene, we considered variants within a 524,288-bp window centered on the transcription start site, corresponding to the genomic interval used for the embedding-based analyses.

We varied both target heritabilities *h*^2^ = *{*0, 0.05, 0.10, 0.25, 0.50, 0.75, 0.95*}* and the proportion of causal variants, *p*_causal_ = *{*0.01, 0.10*}*. For each combination of gene, target heritability, and causal fraction, we generated 100 independent phenotype replicates.

To account for the relationship between allele frequency and genetic effect sizes, we assigned each causal variant *i* an effect-size variance *A*_*i*_ proportional to a power of its allele-frequency variance,

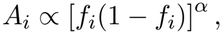

where *f*_*i*_ is the allele frequency of variant *i* and *α* = 0.75 controls the dependence of effect-size variance on allele frequency. For each replicate, the causal effect sizes were then sampled independently as

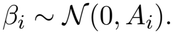

For each replicate, the genetic component was constructed as the sum of the contributions from all causal variants,

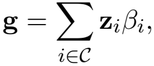

where **z**_*i*_ *∈* ℝ^*n*^ denotes the standardized genotype dosage of variant *i* across the *n* individuals, with mean zero and unit variance.

The phenotype was then generated by adding independent Gaussian residual noise,

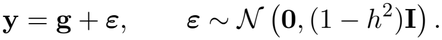

For *h*^2^ = 0, all causal effects were zero and phenotypes consisted entirely of Gaussian noise.

#### Simulations results

For each gene, we compared three variant-binning designs for the genotype-based estimator: 4 bins defined by minor allele count and frequency, and two designs that further split bins by LD score, 6 bins (splitting only the two common-variant bins) and 8 bins (splitting all four). Supplementary Fig. 1 summarizes the HE regression results, showing bias across target values of 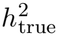, causal fractions, and binning designs (panels a–b), together with the fraction of regressions that converged (panel c). The corresponding REML results are shown in Supplementary Fig. 2. REML had a substantially lower convergence rate and yielded less precise estimates than HE regression; we therefore adopted HE regression as the default estimator throughout the paper. Splitting bins by LD substantially reduced the bias we observed in the 4 bins approach. The two LD-split designs differed mainly in dispersion, which was smaller with 6 bins than with 8 bins. Results were similar for both causal fractions. We therefore adopted the 6-bin design as the default, as this setting yielded the smallest dispersion of estimates around the true value.

### 2.2 Simulation framework for embedding-based heritability estimation

All simulations were performed using real sequence-to-function (S2F) embeddings computed with Borzoi from GTEx donors’ whole-genome sequences, as described in Methods, for 10 randomly selected genes.

We considered two approaches to phenotype simulation. In the first, genetic effects were generated directly from the embedding representation, allowing control over their distribution across embedding dimensions. In the second, genetic effects were simulated from the underlying genotypes, allowing us to assess the extent to which embedding-based heritability estimation recovers genetic variation originating from DNA sequence.

For each simulation setting, we considered target heritabilities *h*^2^ = *{*0, 0.05, 0.10, 0.25, 0.50, 0.75, 0.95*}*. We generated 100 independent simulation for each gene and parameters combination.

For each combination of gene, target heritability, and causal fraction, we generated 100 independent phenotype replicates.

For each gene, we used the standardized embedding matrix **Z** *∈* ℝ^*n×d*^, where *n* is the number of GTEx individuals and *d* is the number of embedding features, to generate genetic effects under the different architectures.

#### Dense assumption

Under the dense assumption, causal effects were distributed across all embedding features. This architecture can be expressed by assigning independent Gaussian effect sizes to the *d* standardized embedding features and constructing the embedding component as

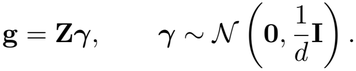

This architecture corresponds to the covariance structure assumed by the embedding-based linear mixed model and therefore provides a baseline for evaluating estimation accuracy under a correctly specified causal architecture.

#### Sparse assumption

Under the sparse assumption, the genetic signal was determined by a single embedding feature, selected randomly for each replicate. Specifically, for a randomly selected feature *j*, the embedding component was defined as

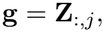

where **Z**_:,*j*_ denotes the values of the *j*-th standardized embedding feature across individuals. This setting represents an architecture in which phenotypic variation depends on a single embedding feature, rather than effects distributed across the full embedding representation.

#### Principal-component-specific assumption

Under the principal-component-specific assumption, the genetic signal was determined by a single embedding principal component, selected randomly for each replicate. We considered only the principal components that explain up to 99% of the embedding variance. Specifically, for a random principal component j, the embedding component was defined as

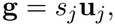

where **u**_*j*_ is the *j*-th column of **U**, obtained from **Z** = **USV**^T^. This design allowed us to investigate how the location of causal effects along principal components explaining different amounts of embedding variance influences heritability estimation, and whether PCA-based dimensionality reduction and whitening affect the recovery of these effects.

#### Phenotype generation

Phenotypes were then constructed as

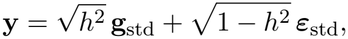

where *h*^2^ denotes the target heritability, **g**_std_ is the standardized embedding component, and ***ε***_std_ is the standardized residual component.

In addition, we evaluated embedding-based heritability estimation using phenotypes simulated directly from genetic variants. In contrast to the embedding-based simulations, where the genetic signal was explicitly constructed from embedding features or principal components, these simulations allowed us to investigate whether embedding representations capture heritable variation originating from the underlying genotype. Genetic effects were simulated as described in Section 2.1, using *p*_*causal*_ = 0.01.

#### Simulation results

For each gene, we compared three embedding representations: the original standardized embedding matrix and two PCA-whitened representations retaining *k* = round(0.7*r*) or *k* = round(0.9*r*) principal components. Supplementary Fig. 3 summarizes the HE regression results, showing bias across target values of 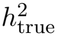, simulation architectures, and embedding representations (panels a–b), together with the fraction of regressions that converged (panel c). The corresponding REML results are shown in Supplementary Fig. 4. Unlike in the genotype-based simulations, REML showed no convergence issues or loss of precision relative to HE in the embedding-based simulations. Nevertheless, we adopted HE for consistency with the genotype-based analyses. The original standardized representation exhibited substantial negative bias under the principal-component-specific architecture and when phenotypes were simulated directly from genetic variants, indicating that it does not consistently support reliable estimation of embedding-based heritability. Both whitened representations substantially reduced this bias. We therefore adopted retention of 70% of the available principal components as the default, as this setting yielded the smallest dispersion of estimates around the true value.

## 3 Supplementary Tables

**Supplementary Table 1.** Borzoi layers used for embedding extraction. Description of the 24 Borzoi layers used for embedding extraction, including layer names, architectural stages, and embedding dimensions.

**Supplementary Table 2.** Genotype-based heritability of whole-blood gene expression. Genotype-based cis-heritability estimates 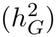 for 16,671 high-quality genes in GTEx whole blood, obtained using a six-component variance model partitioned by allele frequency and linkage disequilibrium.

**Supplementary Table 3.** Embedding-based heritability across Borzoi layers. Embedding-based cis-heritability estimates 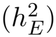 for 500 selected genes across 24 Borzoi layers, comparing the pretrained and randomly initialized models.

**Supplementary Table 4.** Ancestry separability of genotypes and Borzoi embeddings. Ancestry separability (EUR versus AFR) measured by the *R*^2^ of linear models fitted to the first 10 principal components of genotypes and Borzoi embeddings across 500 genes and 24 layers.

**Supplementary Table 5.** Functional enrichment of Borzoi attribution seqlets. Attribution-based functional enrichment results across Borzoi layers for the pretrained and randomly initialized models, including transcription factor and RNA-binding protein motifs, evolutionary conservation (phyloP), and depletion of population variation (gnomAD).

**Supplementary Table 6.** Rare-variant heritability from genotypes and Borzoi embeddings. Cis-heritability estimates for 16,671 high-quality genes in GTEx whole blood, obtained by jointly fitting a rare-variant embedding component (MAF *≤* 0.01) and four genotype-based components partitioned by allele frequency and linkage disequilibrium.

## 4 Supplementary Figures

**Supplementary Figure 1:**
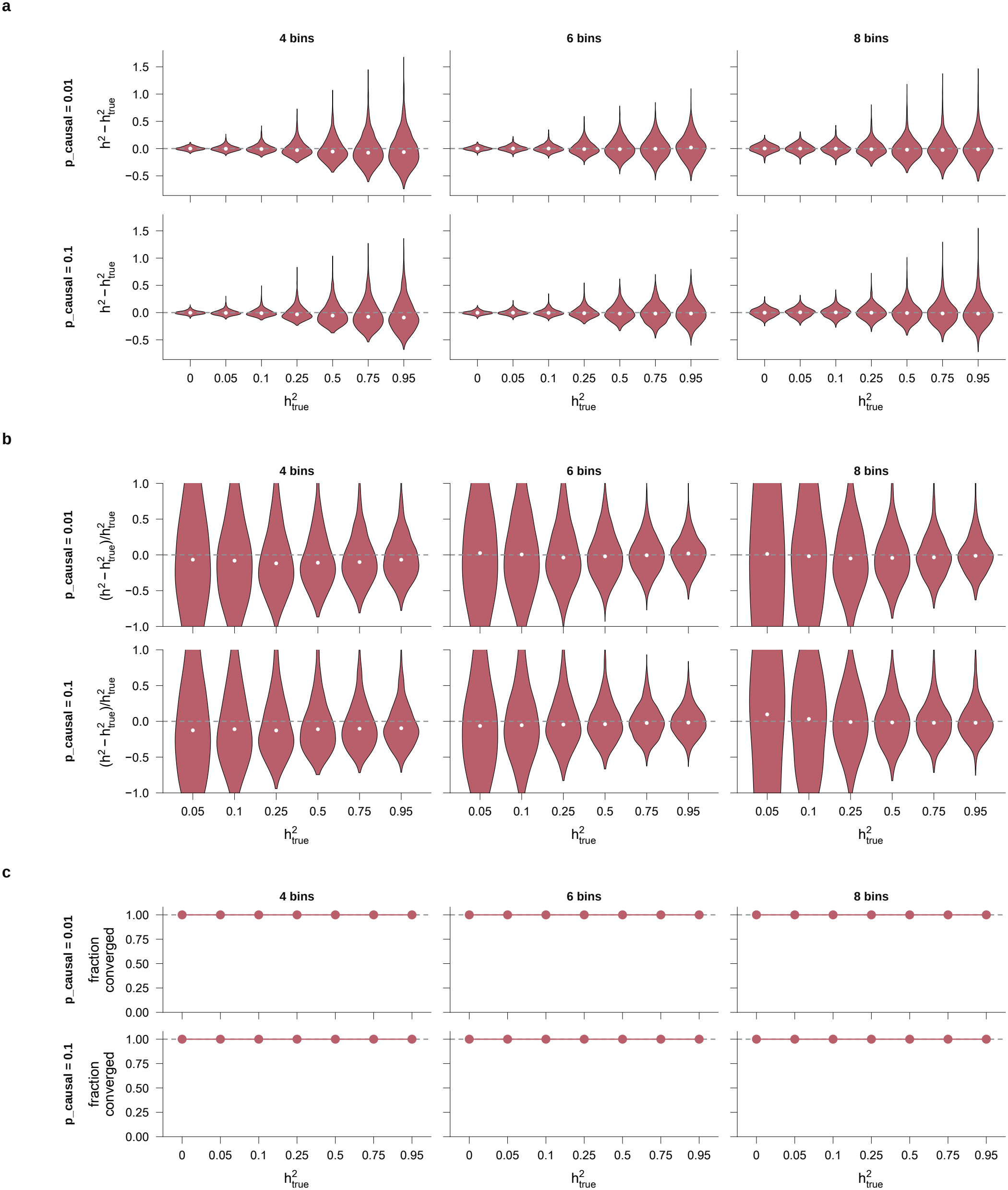
Simulation results showing the bias of the Haseman-Elston (HE) estimator of genotype-based heritability, measured as 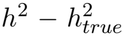 and 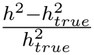 Panel c shows the fraction of regressions that converged. Columns correspond to partitioning schemes with 4, 6, or 8 bins; rows correspond to 1% (top) or 10% (bottom) causal variants. The x-axis shows the true heritability. Each violin shows the distribution of estimation errors across 100 simulation replicates for 10 random genes. White dots indicate medians. Further details of the simulation experiments are explained in Supplementary Note 2.1.

**Supplementary Figure 2:**
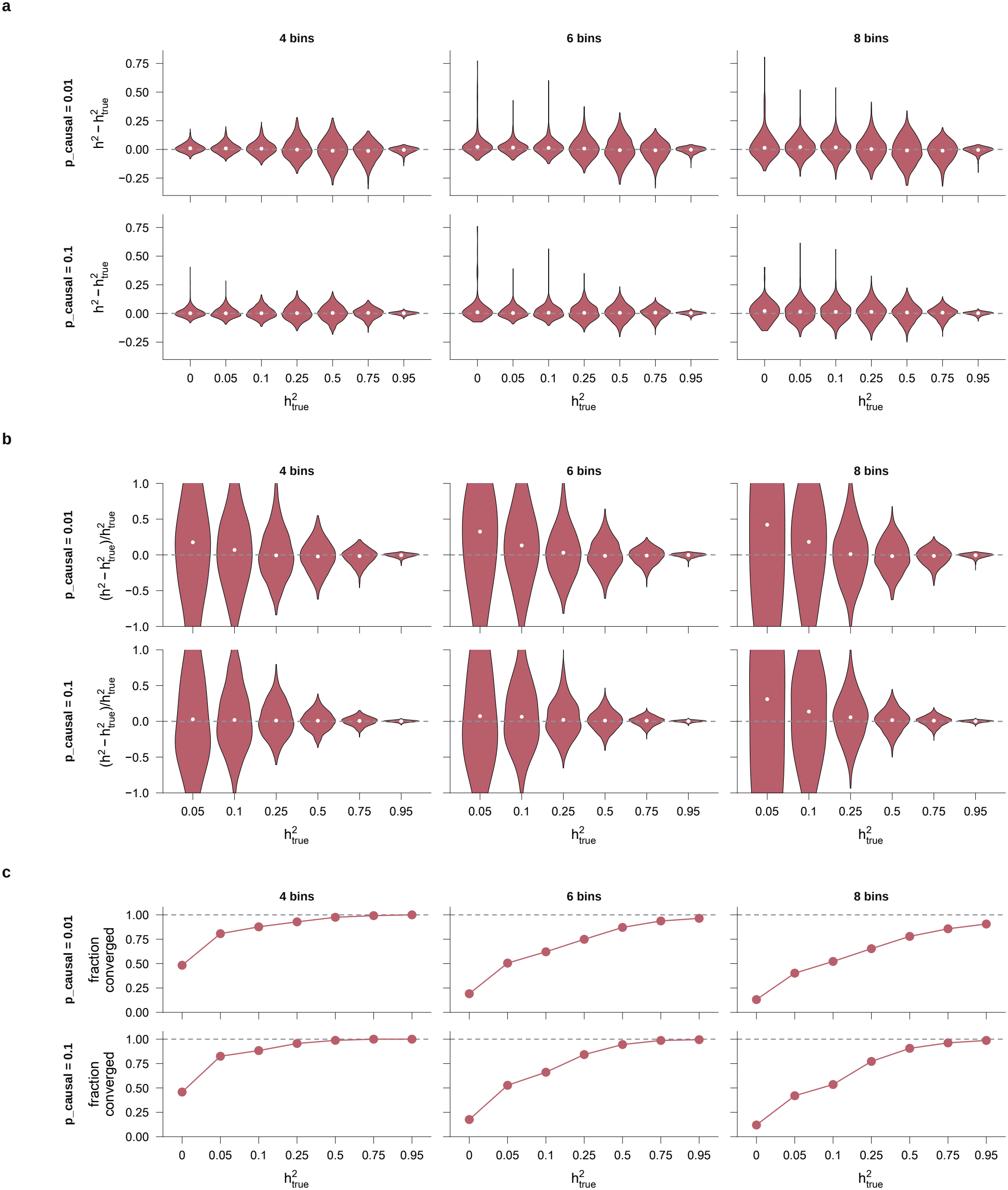
Simulation results showing the bias of the REML estimator of genotype-based heritability, measured as 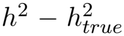 and 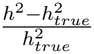. Panel c shows the fraction of regressions that converged. Columns correspond to partitioning schemes with 4, 6, or 8 bins; rows correspond to 1% (top) or 10% (bottom) causal variants. The x-axis shows the true heritability. Each violin shows the distribution of estimation errors across 100 simulation replicates for 10 random genes. White dots indicate medians. Further details of the simulation experiments are explained in Supplementary Note 2.1.

**Supplementary Figure 3:**
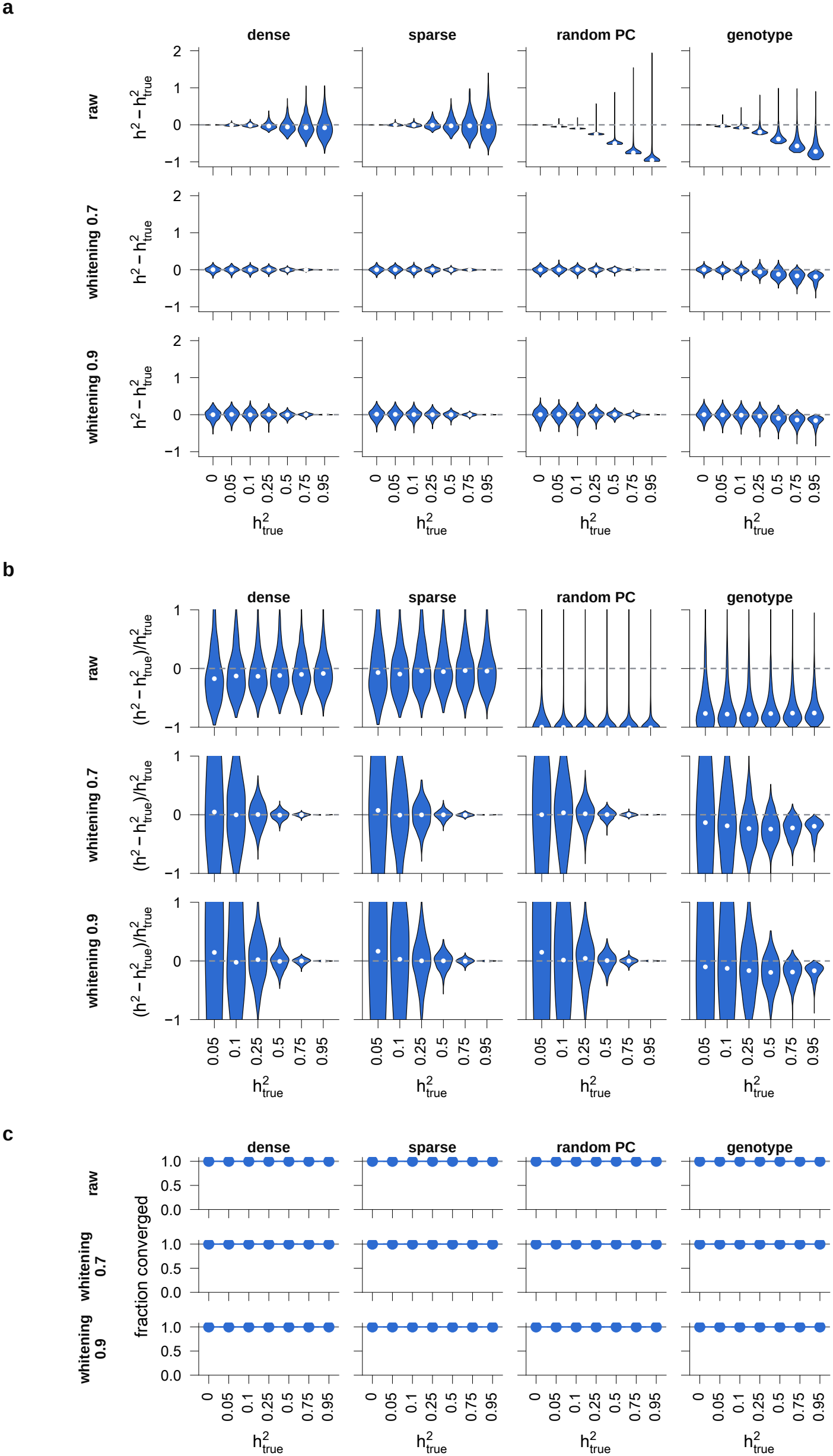
Simulation results showing the bias of the Haseman–Elston (HE) estimator of embedding-based heritability, measured as 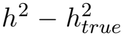 and 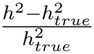. Panel c shows the fraction of regressions that converged. Columns correspond to different assumptions of feature effect size distribution; rows correspond to different embedding representations. The x-axis shows the true heritability. Each violin shows the distribution of estimation errors across 100 simulation replicates for 10 random genes. White dots indicate medians. Further details of the simulation experiments are explained in Supplementary Note 2.2.

**Supplementary Figure 4:**
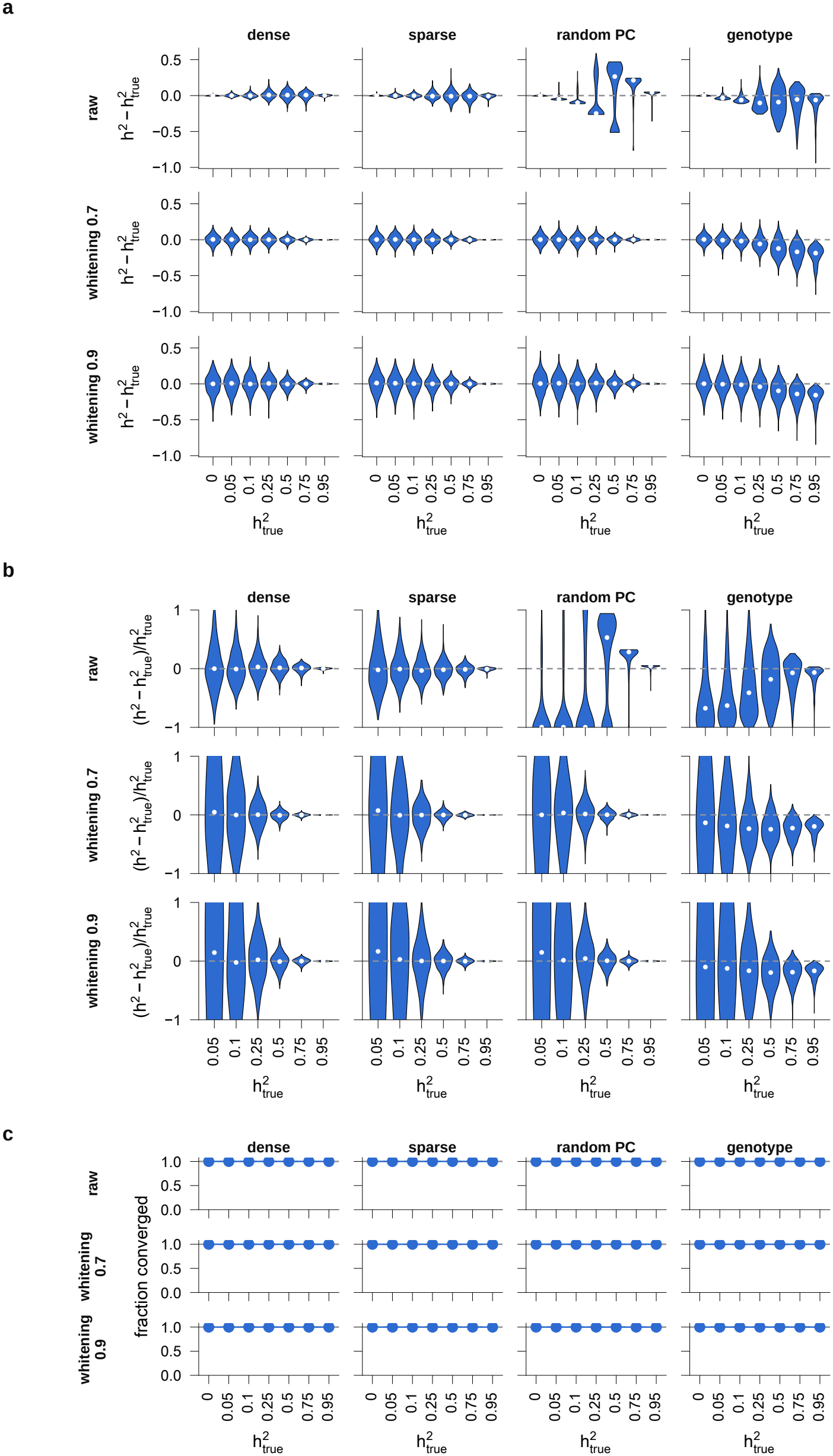
Simulation results showing the bias of the REML estimator of embedding-based heritability, measured as 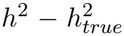 and 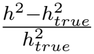. Panel c shows the fraction of regressions that converged. Columns correspond to different assumptions of feature effect size distribution; rows correspond to different embedding representations. The x-axis shows the true heritability. Each violin shows the distribution of estimation errors across 100 simulation replicates for 10 random genes. White dots indicate medians. Further details of the simulation experiments are explained in Supplementary Note 2.2.

**Supplementary Figure 5:**
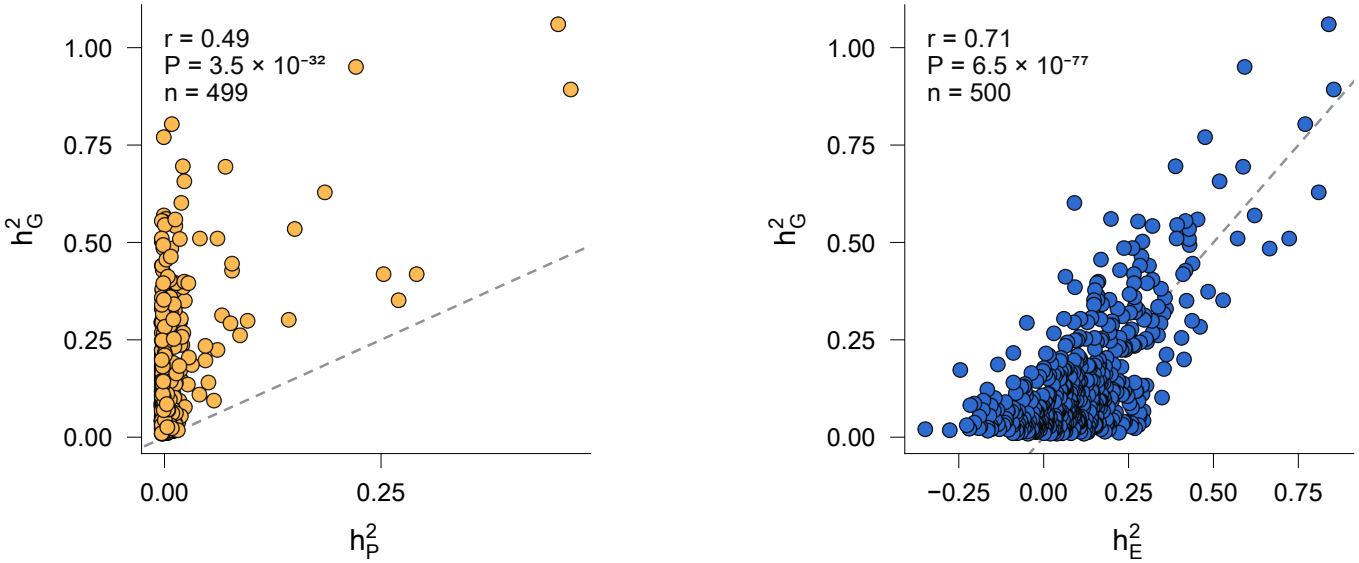
Analyses use GEUVADIS lymphoblastoid cell-line (n = 449 donors) and 500 randomly selected genes with 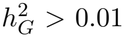. Genotype-based expression heritability 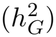 plotted against the heritability explained by Borzoi’s whole-blood predictions (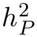, left) and by embeddings from the penultimate layer (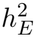, right). Each point represents a gene; dashed lines indicate equality (*y* = *x*); r, Pearson correlation.

**Supplementary Figure 6:**
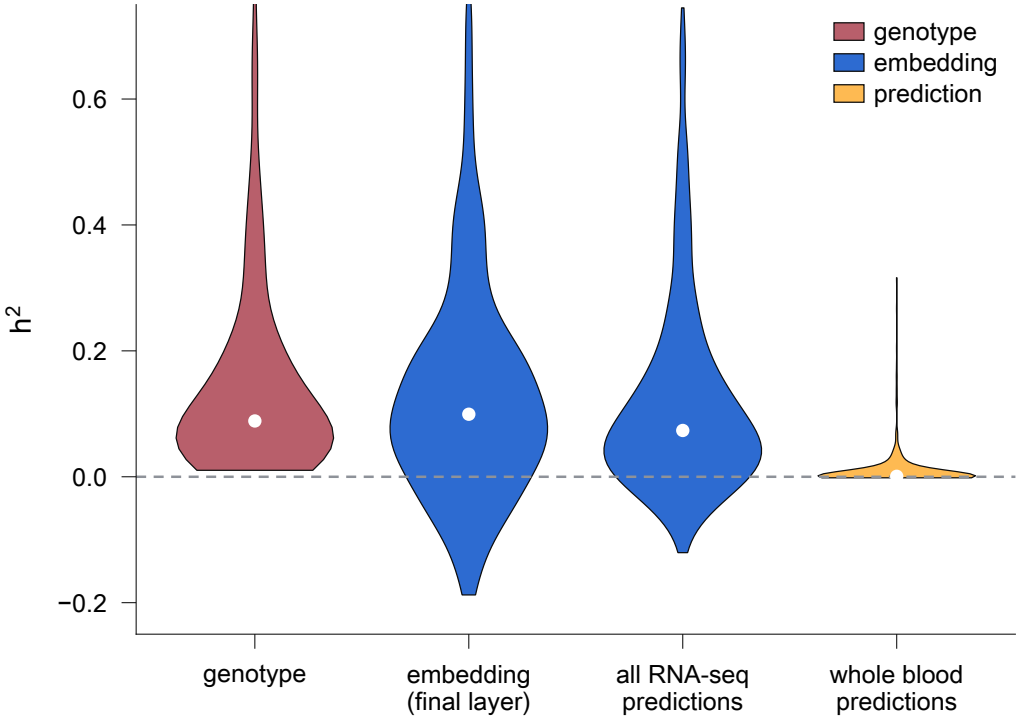
Distribution of heritability across 500 random genes. From left to right: geno-type heritability; embedding heritability, computed from the final-layer of Borzoi (Methods); heritability explained by all Borzoi RNA-seq predictions; heritability explained by Borzoi whole-blood predictions. The white dot is the median.

**Supplementary Figure 7:**
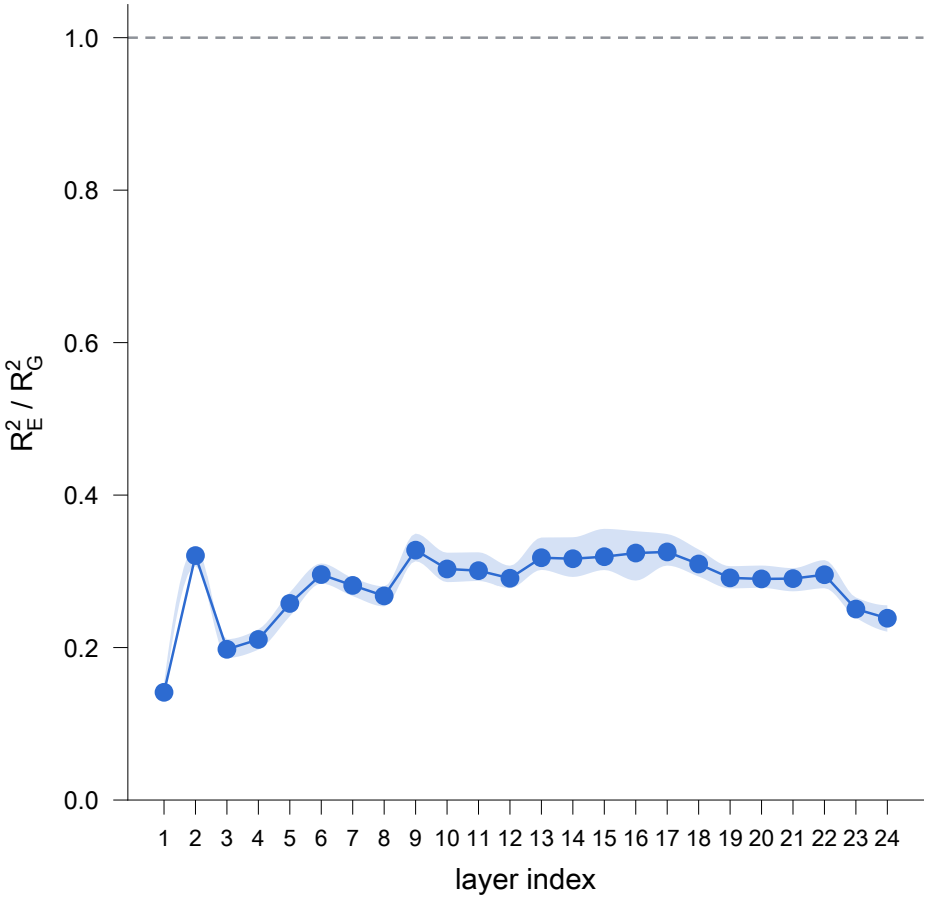
Ratio between the ancestral variance explained by the first ten principal components of embeddings across Borzoi layers, compared with local-genotype-derived first ten principal components. Variance explained is measured with *R*^2^. The blue line represents the median of the per-gene R2 ratio across the genes and the shaded band is the interquartile range (25th to 75th percentile) of the per-gene ratio. The horizontal red shaded line represents a ratio of 1. Results are shown across Borzoi layers.

**Supplementary Figure 8:**
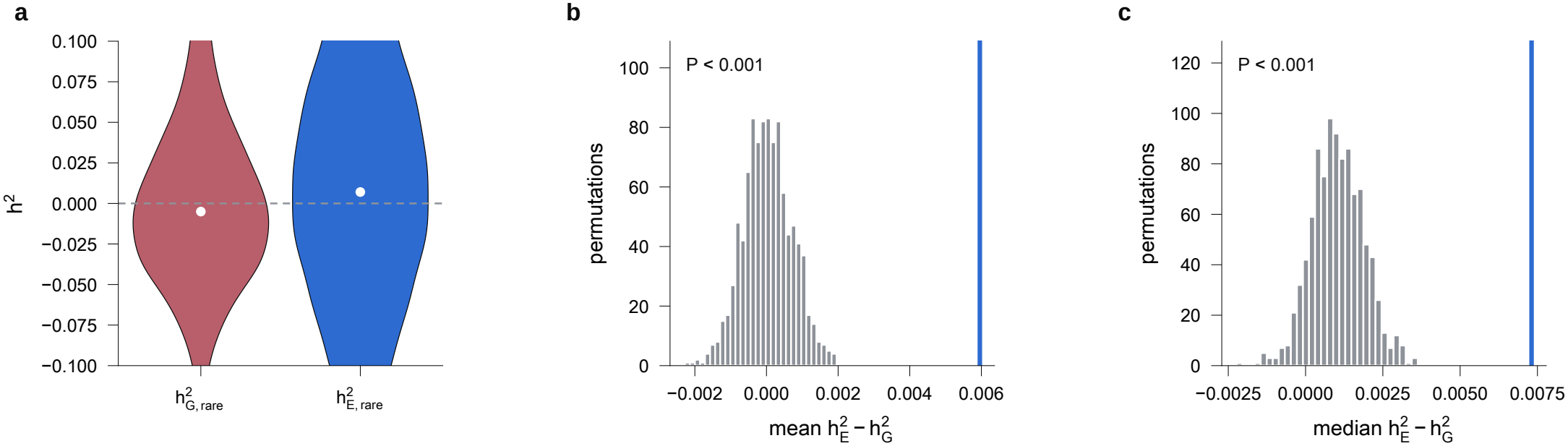
a, Distribution across genes of the rare-variant heritability estimated from the genotype and from Borzoi embeddings. The white dot is the median and the dashed line marks zero. b, c, Permutation test of 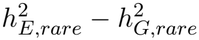. The phenotype was shuffled across donors and the difference was re-estimated for each shuffle. Each histogram shows, across shuffles, the mean (b) and median (c) of this difference across genes. The blue line is the observed value.

## REFERENCES

1. Gamazon, E. R. et al. A gene-based association method for mapping traits using reference transcriptome data. Nat. Genet. 47, 1091–1098 (2015).

2. Wheeler, H. E. et al. Survey of the heritability and sparse architecture of gene expression traits across human tissues. PLoS Genet. 12, e1006423 (2016).

3. Gusev, A. et al. Integrative approaches for large-scale transcriptome-wide association studies. Nat. Genet. 48, 245–252 (2016).

4. Avsec, Ž. et al. Effective gene expression prediction from sequence by integrating long-range interactions. Nat. Methods 18, 1196–1203 (2021).

5. Linder, J., Srivastava, D., Yuan, H., Agarwal, V. & Kelley, D. R. Predicting RNA-seq coverage from DNA sequence as a unifying model of gene regulation. Nat. Genet. 1–13 (2025).

6. Avsec, Ž. et al. Advancing regulatory variant effect prediction with AlphaGenome. Nature 649, 1206–1218 (2026).

7. Clarke, B. et al. Integration of variant annotations using deep set networks boosts rare variant association testing. Nat. Genet. 56, 2271–2280 (2024).

8. Hernandez, R. D. et al. Ultrarare variants drive substantial cis heritability of human gene expression. Nat. Genet. 51, 1349–1355 (2019).

9. Pampari, A. et al. ChromBPNet: bias factorized, base-resolution deep learning models of chromatin accessibility reveal cis-regulatory sequence syntax, transcription factor footprints and regulatory variants. bioRxivorg 2024.12.25.630221 (2025) doi:10.1101/2024.12.25.630221.

10. Jaganathan, K. et al. A unified framework for quantitative splicing and transcript prediction. bioRxivorg 2026.10.08.757673 (2026) doi:10.64898/2026.10.08.757673.

11. Zeng, T. & Li, Y. I. Predicting RNA splicing from DNA sequence using Pangolin. Genome Biol. 23, 103 (2022).

12. Jaganathan, K. et al. Predicting Splicing from Primary Sequence with Deep Learning. Cell 176, 535–548.e24 (2019).

13. Xu, C. et al. Reference-informed prediction of alternative splicing and splicing-altering mutations from sequences. Genome Res. 34, 1052–1065 (2024).

14. Sasse, A. et al. Benchmarking of deep neural networks for predicting personal gene expression from DNA sequence highlights shortcomings. Nat. Genet. 55, 2060–2064 (2023).

15. Tu, X., Spiro, A., Chikina, M. & Mostafavi, S. A modality gap in personal-genome prediction by sequence-to-function models. bioRxiv 2026.02.01.702969 (2026) doi:10.64898/2026.02.01.702969.

16. Spiro, A. E. et al. A scalable approach to investigating sequence-to-function predictions from personal genomes. Nat. Methods 23, 1308–1312 (2026).

17. Drusinsky, S., Whalen, S. & Pollard, K. S. Deep-learning prediction of gene expression from personal genomes. Genome Biol. 27, 19 (2026).

18. Yang, J. et al. Common SNPs explain a large proportion of the heritability for human height. Nat. Genet. 42, 565–569 (2010).

19. Speed, D., Hemani, G., Johnson, M. R. & Balding, D. J. Improved heritability estimation from genome-wide SNPs. Am. J. Hum. Genet. 91, 1011–1021 (2012).

20. Evans, L. M. et al. Comparison of methods that use whole genome data to estimate the heritability and genetic architecture of complex traits. Nat. Genet. 50, 737–745 (2018).

21. Rastogi, R., Reddy, A. J., Chung, R. & Ioannidis, N. M. Fine-tuning sequence-to-expression models on personal genome and transcriptome data. Genome Biol. 27, 234 (2026).

22. Huang, C. et al. Personal transcriptome variation is poorly explained by current genomic deep learning models. Nat. Genet. 55, 2056–2059 (2023).

23. Drusinsky, S. & Pollard, K. S. Causal variant underestimation is a major overlooked driver of sequence-to-function model underperformance. Genetics 2026.09.08.750172 (2026) doi:10.64898/2026.09.08.750172.

24. Shrikumar, A., Greenside, P. & Kundaje, A. Learning Important Features Through Propagating Activation Differences. in International Conference on Machine Learning 3145–3153 (PMLR, 2017).

25. Keinan, A. & Clark, A. G. Recent explosive human population growth has resulted in an excess of rare genetic variants. Science 336, 740–743 (2012).

26. Tennessen, J. A. et al. Evolution and functional impact of rare coding variation from deep sequencing of human exomes. Science 337, 64–69 (2012).

27. Kingdom, R. et al. Rare genetic variants in genes and loci linked to dominant monogenic developmental disorders cause milder related phenotypes in the general population. Am. J. Hum. Genet. 109, 1308–1316 (2022).

28. Forrest, I. S. et al. Population-Based Penetrance of Deleterious Clinical Variants. JAMA 327, 350–359 (2022).

29. Castel, S. E. et al. Modified penetrance of coding variants by cis-regulatory variation contributes to disease risk. Nat. Genet. 50, 1327–1334 (2018).

30. Pazokitoroudi, A. et al. Efficient variance components analysis across millions of genomes. Nat. Commun. 11, 4020 (2020).

31. Liu, Z. et al. Comprehensive gene heritability estimation reveals the genetic architecture of rare coding variants underlying complex traits. bioRxivorg 2025.10.07.681018 (2025) doi:10.1101/2025.10.07.681018.

32. Grant, C. E., Bailey, T. L. & Noble, W. S. FIMO: scanning for occurrences of a given motif. Bioinformatics 27, 1017–1018 (2011).

33. Vorontsov, I. E. et al. HOCOMOCO in 2024: a rebuild of the curated collection of binding models for human and mouse transcription factors. Nucleic Acids Res. 52, D154–D163 (2024).

34. Rosado-Tristani, D. A. et al. CisBP-RNA: a web resource for eukaryotic RNA-binding proteins and their motifs. Nucleic Acids Res. 54, D98–D105 (2026).

35. Pazokitoroudi, A. et al. A scalable and robust variance components method reveals insights into the architecture of gene-environment interactions underlying complex traits. Am. J. Hum. Genet. 111, 1462–1480 (2024).

36. Chang, C. C. et al. Second-generation PLINK: rising to the challenge of larger and richer datasets. Gigascience 4, 7 (2015).

37. Danecek, P. et al. Twelve years of SAMtools and BCFtools. Gigascience 10, giab008 (2021).

38. Hingerl, J. C., Karollus, A. & Gagneur, J. Flashzoi: an enhanced Borzoi for accelerated genomic analysis. Bioinformatics 41, btaf467 (2025).

39. Kokhlikyan, N. et al. Captum: A unified and generic model interpretability library for PyTorch. arXiv [cs.LG] (2020) doi:10.48550/arXiv.2009.07896.

40. Schreiber, J. Tangermeme: a toolkit for understanding cis-regulatory logic using deep learning models. Nat. Methods 23, 2136–2140 (2026).

41. Lal, A. et al. Decoding sequence determinants of gene expression in diverse cellular and disease states. Nat. Methods 23, 1138–1151 (2026).

42. Lal, A., Gunsalus, L., Nair, S., Biancalani, T. & Eraslan, G. gReLU: a comprehensive framework for DNA sequence modeling and design. Nat. Methods 22, 2253–2257 (2025).

43. Di Tommaso, P. et al. Nextflow enables reproducible computational workflows. Nat. Biotechnol. 35, 316–319 (2017).

44. Köster, J. & Rahmann, S. Snakemake--a scalable bioinformatics workflow engine. Bioinformatics 28, 2520–2522 (2012).

## Bibliography

[1] Mathew, B., Léon, J. & Sillanpää, M. J. A novel linkage-disequilibrium corrected genomic relationship matrix for SNP-heritability estimation and genomic prediction. Heredity 120, 356–368 (2018).

[2] Ma, R. & Dicker, L. H. The Mahalanobis kernel for heritability estimation in genome-wide association studies: fixed-effects and random-effects methods. arXiv preprint arXiv:1901.02936 (2019). URL https://arxiv.org/abs/1901.02936.

